# Cognate amino acid therapies provide preclinical benefit in 19 *C. elegans* models of ARS2 deficiency

**DOI:** 10.64898/2026.02.03.703549

**Authors:** Cristina Remes, Kelsey Keith, Rui Xiao, Vernon E. Anderson, Donna M. Iadarola, Eiko Nakamaru-Ogiso, Neal D. Mathew, Marni J. Falk

## Abstract

Mitochondrial aminoacyl-tRNA synthetases (mt-ARS) are essential mitochondrial translation machinery components that catalyze mitochondrial transfer RNAs (tRNAs) charging with their cognate amino acid. Although mt-ARS have a common biochemical function, patients with mt-ARS pathogenic variants commonly develop neurological disorders with varying phenotypes, severity spectrum, and age of onset. Cognate amino acid supplementation has shown reported benefits in select cases of both mt-ARS (*ARS2*) and cytosolic (*ARS1*) deficiencies, although the safety and potential benefits of this candidate therapy approach across the full spectrum of mt-ARS disorders remain unclear. Here, *C. elegans* models were systematically generated for all 19 mitochondrial mt-ARS genes by feeding RNAi knockdown for one or two generations. mt-ARS deficient animals at baseline and upon cognate amino acid treatment were studied at the level of linear growth, neuromuscular activity, lifespan, mitochondrial physiology, and fertility. Results demonstrated that cognate amino acid treatment in a dose-dependent fashion consistently improved worm linear growth and neuromuscular activity, and reduced mitochondrial unfolded protein response stress, in all 19 knockdown models. It further rescued impaired fertility of *hars-1* and *fars-2* knockdown strains. Collectively, these preclinical studies provide compelling evidence to warrant future cognate amino acid treatment study in rigorous clinical trials spanning all human mt-ARS deficiencies.

## INTRODUCTION

Primary mitochondrial diseases (PMD) share a final common pathway of impaired oxidative phosphorylation (OXPHOS) capacity due to pathogenic variants in a wide range of nuclear or mitochondrial DNA genes. Among these, genes that support the process of mitochondrial translation are essential to synthesize 13 mtDNA-encoded respiratory chain (RC) core subunits of OXPHOS complexes I, III, IV, and V. Impaired mitochondrial translation may result from pathogenic variants in genes encoding 22 mitochondrial transfer RNAs (mt-tRNAs), 2 mitochondrial ribosomal RNAs (mt-rRNAs), nuclear-encoded mitochondrial ribosomal proteins, or 19 nuclear-encoded mitochondrial aminoacyl-tRNA synthetases (mt-ARS, encoded by genes generally delineated by each amino acid symbol followed by ‘ARS2’) ^1–3^. mt-ARS proteins are highly evolutionarily-conserved enzymes that catalyze charging of mt-tRNAs with their cognate (specific) amino acid, a process that ensures the translational accuracy of mitochondrial protein synthesis. As an exemplar, alanine mt-ARS (mt-AARS) specifically charges tRNA^Ala^ with its cognate amino acid, alanine. Defects in the mt-ARS proteins directly impair mitochondrial translation, OXPHOS capacity, and cellular energy production. Most mt-ARS disorders are inherited in an autosomal recessive although GARS2 heterozygous pathogenic gain of function variants cause autosomal dominant Charcot-Marie Tooth disease type 2D ^4^. mt-ARS deficiencies commonly manifest in the central nervous system (CNS), including 8 that lead to encephalopathies and 4 to leukodystrophies, although clinical phenotypes are often heterogeneous and tissue-specific ^5,6^. mt-AARS, mt-GARS, and mt-LARS deficiencies may cause cardiomyopathies ^7–9^; mt-HARS and mt-LARS deficiencies cause Perrault syndrome, characterized by ovarian dysgenesis and hearing loss ^5,10^; and additional phenotypes may include isolated hearing loss (mt-MARS, mt-NARS), intellectual disability (mt-RARS, mt-WARS), Myopathy, Lactic Acidosis and Sideroblastic Anemia (MLASA) syndrome (mt-YARS), and Hyperuricemia, Pulmonary hypertension, Renal failure in infancy, and Alkalosis (HUPRA) syndrome (mt-SARS)^5^.

*mt-ARS* proteins are mainly although not exclusively encoded by nuclear genes whose naming convention begins with their amino acid code followed by “ARS2”, translated in the cytoplasm, and subsequently imported into mitochondria. Each of the 19 mt-tRNAs in human mitochondria are charged by a specific mt-ARS protein, with the exception of mt-tRNA^Gln^ for glutamine. *In vitro* ^11^ and *in vivo* ^12^ studies demonstrated that mt-tRNA^Gln^ is formed by transamidation of the mischarged Glu-tRNA^Gln^ in mammalian mitochondria. Furthermore, in humans only 2 ARS proteins (GARS1 and KARS1) are functional in translation both within the cytosol and mitochondria ^13^; all other mt-ARS are functional only within the mitochondria. In *C. elegans*, 5 mt-ARS proteins are functional in both cytosol and mitochondria: *cars-1*, gars-1, *hars-1*, *kars-1,* and *tars-1* ^14^ Moreover, *aars-1* in *C. elegans* has been shown to localize to the mitochondria, while *aars-2* is localized in the cytosol ^15^. In addition to their aminoacylation activity, evidence suggests that mt-ARS proteins may have non-canonical functions, participating in angiogenesis ^16^ and integrated stress response activation ^17^. Similar ‘moonlighting’ functions have been previously described for cytoplasmic tRNA synthetases ^18–20^. Moreover, ARS2-deficient neuronal models exhibit different compensatory mechanisms, which may contribute to variable clinical phenotypes and tissues affected in ARS2 models ^21^.

Currently, no FDA-approved therapies or cures exist for mt-ARS deficiencies. Preclinical modeling in fibroblasts derived from ARS1 patients has shown that fibroblasts from ARS1 patients are sensitive to limited cognate amino acid availability, and that oral supplementation with cognate amino acids has a beneficial effect for several different disorders caused by deficiencies of cytoplasmic ARS genes.^22^ However, the exact mechanism, efficacy, dosing, formulation, and safety of cognate amino acid supplementation in ARS2 disorders remain poorly understood. Systematic preclinical animal model analysis of the physiologic consequences and response to cognate amino acid supplementation as a possible treatment strategy has not previously been reported for the mt-ARS class of genetic disease. Here, we first characterized the phenotypic and biochemical effects for all 19 conserved mt-ARS genes in *C. elegans* by individually knocking-down the mRNA for each mt-ARS to reduce the amount of each enzyme using feeding RNA interference (RNAi) technology ^23^. Cognate amino acid treatment effects at different doses were then modeled to determine their tolerability and phenotypic rescue in all 19 mt-ARS knockdown worm strains, with particular focus on *dars-2(RNAi)* knockdown worms.

## RESULTS

### 19 mt-ARS *C. elegans* knockdown strains generated by feeding RNAi for one generation consistently manifest growth impairment with variable effects on neuromuscular activity, lifespan, and fertility

High sequence similarity exists between *C. elegans* and human mt-ARS proteins, ranging from 36% (CARS1) to 75% (KARS1) (Supplementary Table 1). Therefore, worms offer a highly informative animal model to enable the preclinical study of the *in vivo* functional effects of mt-ARS proteins’ deficiencies on quantifiable physiologic and biochemical outcomes. 19 mt-ARS knockdown strains were generated by feeding *C. elegans* beginning at the embryo stage with 2 mM IPTG and RNAi bacteria specific for each mt-ARS gene (Supplementary Table 1). A significant decrease in worm length was consistently observed relative to wild-type control (N2 Bristol worms fed empty vector *L4440* bacteria) (Figure 1A), ranging from 10% to 38% depending on the specific gene, with greatest degree of growth impairment in *hars-1(RNAi)*, *cars-1(RNAi)*, and *kars-1(RNAi)* knockdown worms. These data demonstrate that a linear growth defect is a common phenotype of mt-ARS deficiency in worms.

**Figure 1.**
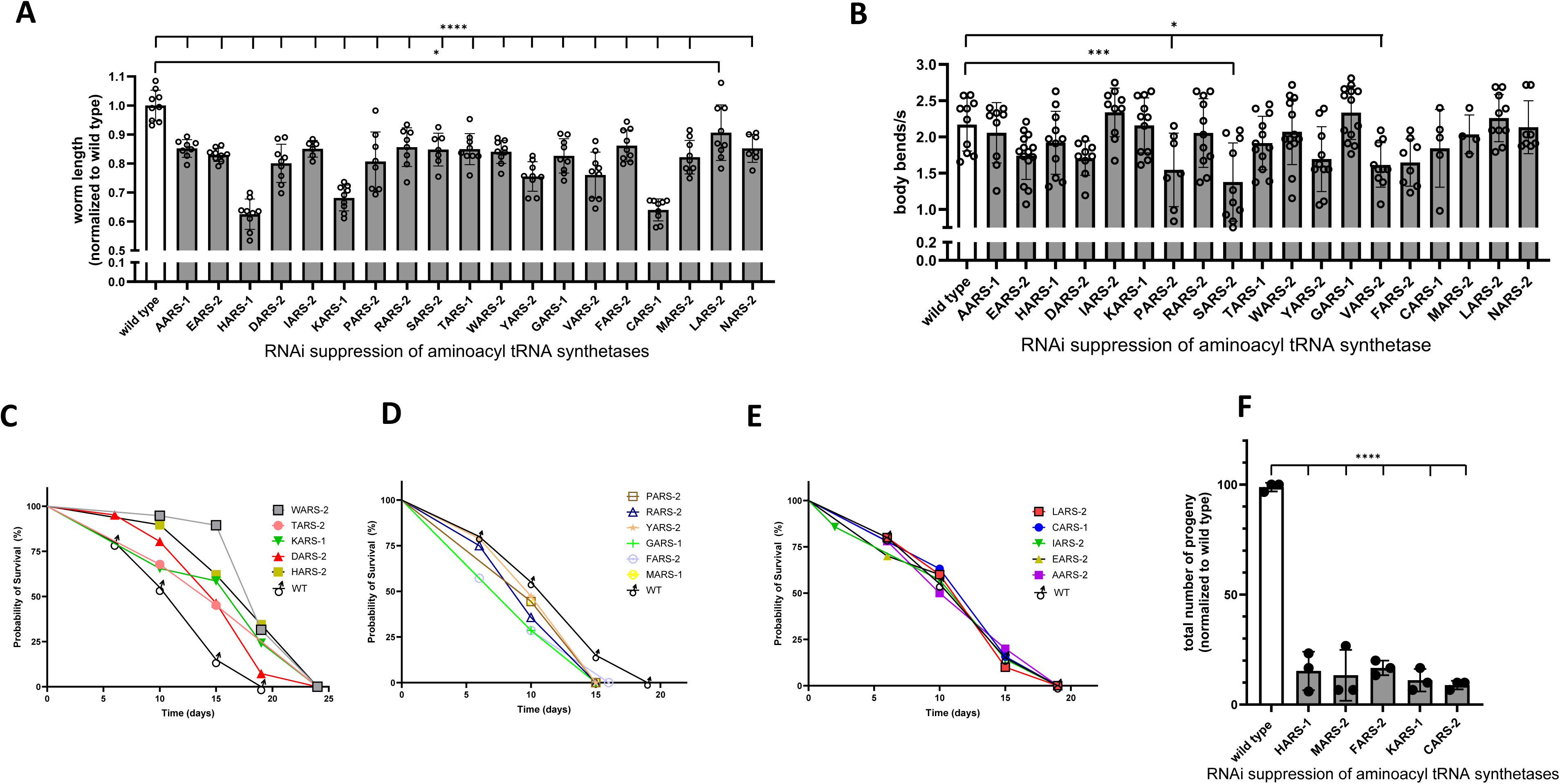
Phenotypic characterization of animal health in first-generation feeding RNAi knockdown of 19 mt-ARS *C. elegans* strains. In all figures, gray bars convey RNAi knockdown of the indicated mt-ARS. **(A)** Worm linear growth analysis shows a consistent decrease in worm length across the nineteen mt-ARS RNAi knockdown worms relative to the empty vector control. Data represent the mean ± SD of 10 individual animals. Variation was detected by two-sided, 1-way ANOVA with Dunnet’s post hoc multiple comparisons test. *: p <0.05; ****: p<0.0001 **(B)** Worm neuromuscular activity assessed showed decreased swimming-based thrashing activity for a set of the knockdown worms identified 1-way ANOVA test. Data represent the mean ± SD of 20 individual animals. **(C-E)** Lifespan analysis of the mt-ARS knockdown worms shows distinct changes in lifespan compared to wild type, with some strains showing (C) increased lifespan (D) decreased lifespan and (E) no significant change in lifespan. **(F)** Five mt-ARS knockdown strains showed a dramatic decrease (∼85-90%) in progeny count compared to wild type animals. Data represent the mean ± SD of 3 individual animals. Variation was detected by Unpaired t test with Welch’s correction. *: p <0.05, **: p<0.01, ***: p<0.001, ****: p<0.0001

Since most mt-ARS disorders impair neurologic and/or muscular function, we characterized neuromuscular activity in the mt-ARS knockdown worms using a thrashing assay of worms placed in liquid media ^24,25^. Interestingly, only a few of the mt-ARS knockdown strains exposed to RNAi for one generation showed significantly decreased worm activity (measured as body bends per second) relative to the wild-type control (Figure 1B). Among these, *dars-2*, *vars-2, yars-2* and *sars-2* deficiencies yielded the largest decrease (∼20%) in worm thrashing activity.

Given the relatively short lifespan of 2 weeks on average and ∼4 week maximal at 20°C of wild-type (N2 Bristol) *C. elegans*, we investigated the effect of mt-ARS knockdown on worm survival by interval WormScan analysis ^26^. Interestingly variable effects were seen, where one subset of mt-ARS knockdown strains had no lifespan alteration (Figure 1C), while the lifespan of other subsets was significantly increased (Figure 1D) or decreased (Figure 1E) relative to wild-type worms grown at 20°C. Thus, survival was not a consistently impaired outcome of one generation of mt-ARS knockdown across all 19 strains.

Since mitochondrial translation defects have been reported to reduce fertility in humans ^27^, we analyzed fertility of the knockdown strains using a standard *C. elegans* progeny count assay. Results showed that RNAi knockdown for one generation of *hars-1, mars-2*, *fars-2*, *kars-1* and *cars-1* resulted in a significant decrease (by 84%, 87%, 83%, 88% and 91%, respectively) in progeny numbers compared to wild-type worms (Figure 1D).

Collectively, these data demonstrate that mt-ARS protein deficiencies induced by exposure to continuous feeding RNAi for one generation consistently induced impairment of animal growth but had variable effects on neuromuscular activity, lifespan, and fertility (progeny).

### 19 mt-ARS *C. elegans* knockdown strains generated by feeding RNAi for one generation manifested variable *in vivo* induction of mitochondrial stress and reduction of mitochondrial membrane potential

Given the critical role of mitochondrial translation for OXPHOS function, high-content imaging analysis (CX5, ThermoFisher) was used to quantify the effect of mt-ARS knockdown on mitochondrial unfolded protein response (UPR^mt^) stress induction, which is a common physiologic response to impaired OXPHOS capacity ^28–30^. Specifically, we used a *C. elegans* transgenic line containing two fluorescent reporters (*hsp-6p::gfp* and *myo-2p::mCherry)*, where mCherry fluorescence localized in the worm head to normalize worm number per well and GFP fluorescence indicated the degree of mitochondrial unfolded protein stress response induction ^31^. Worms were grown on solid media throughout development until reaching 1 day of adulthood past the fourth larval stage (L4 + 1 day), when they were transferred to a 384-well plate for fluorescence quantification. Interestingly, highly significant UPR^mt^ stress induction levels were observed for each of the 19 mt-ARS knockdown strains (Figure 2A, 2B). The strongest degree of UPR^mt^ stress response induction was seen in *dars-2(RNAi)* knockdown worms, which had an approximately 10-fold higher *hsp-6p*::GFP fluorescence induction relative to other mt-ARS knockdown strains following grown on RNAi for one generation (Figure 2B).

**Figure 2.**
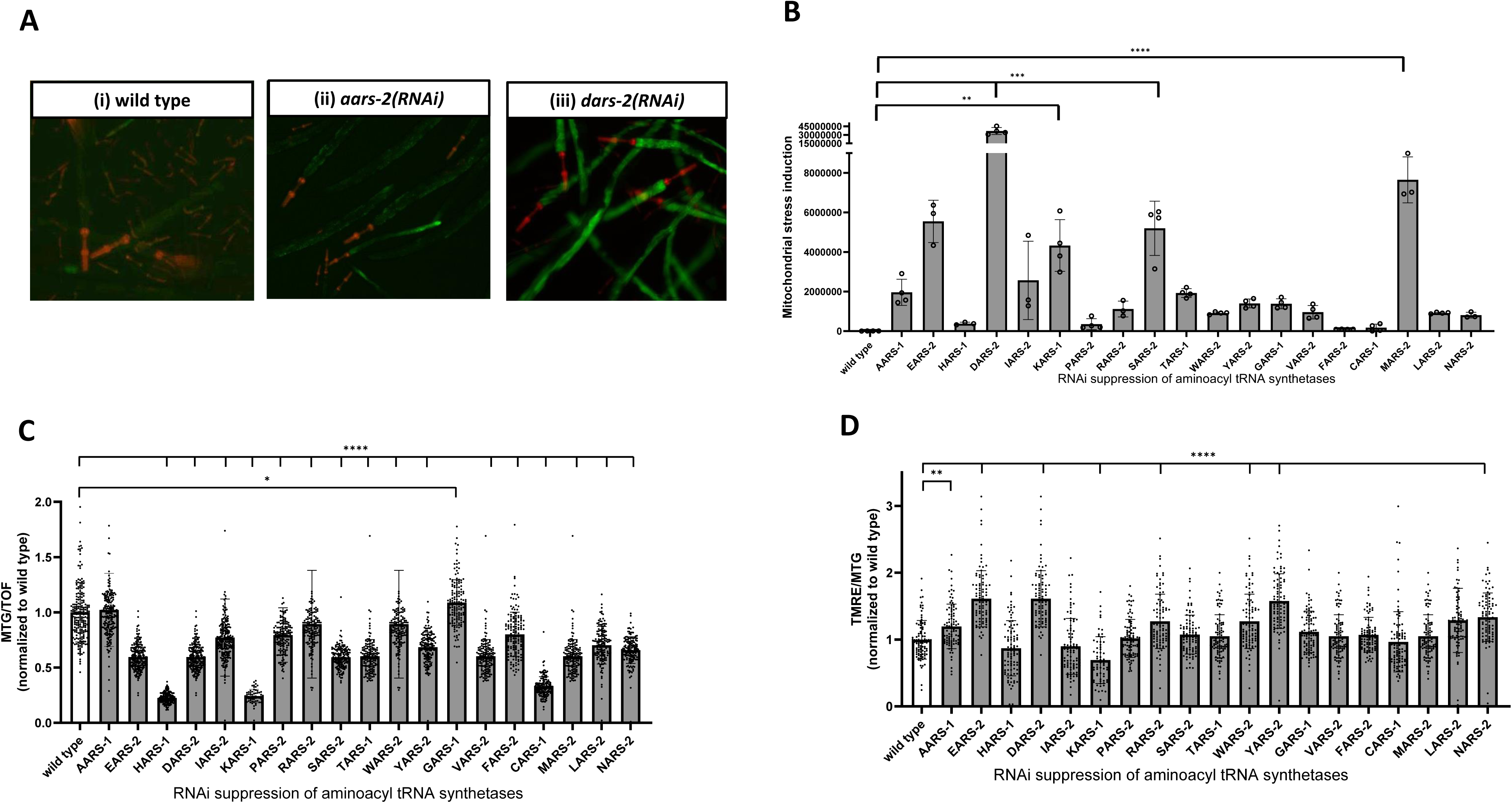
Phenotypic characterization of mitochondrial physiology in the first-generation of mt-ARS knockdown worms. **(A)** High-content imaging of two RNAi knockdown strains showing distinct responses in UPR^mt^ stress induction upon mt-ARS knockdown: large increase in UPR^mt^ stress in *dars-2(RNAi*) and small increase in UPR^mt^ stress in *aars-1(RNAi)* compared to wild type. Shown are representative images of 3 biological replicates. **(B)** Quantification of UPR^mt^ stress for the full set of ARS RNAi worms, showing distinct induction of UPR^mt^ stress induction upon knockdown of mt-ARS genes. Data represent the mean ± SD of 10 individual wells containing ∼50 worms. Variation was detected by 1-way ANOVA test. **(C)** Quantification of mitochondrial mass after MitoTracker Green (MTG) labeling in the mt-ARS knockdown worms, showing distinct responses in mitochondrial content. For each condition, approximately 300 worms were studied in three biological replicates. Variation was detected by 1-way ANOVA test. **(D)** Quantification of the relative mitochondrial membrane potential in the mt-ARS knockdown worms, assessed by normalizing TMRE to MTG florescence. Approximately 300 animals were measured for each condition, in three biological replicates. Variation was detected by 1-way ANOVA test. *: p <0.05, **: p<0.01, ***: p<0.001, ****: p<0.0001

*In vivo* analysis of mitochondrial content and integrated respiratory capacity at the level of mitochondrial membrane potential was relatively quantified in each mt-ARS deficient strain by fluorescence microscopy with MitoTracker^®^ green (MTG) and tetramethylrhodamine, ethyl ester (TMRE), respectively. All mt-ARS deficient strains, except for *aars-1*, showed significant reduction following one generation of RNAi exposure in mitochondrial content (MTG) as normalized to the size of each animal (time of flight) (Figure 2C). The greatest degree of mitochondrial depletion was seen in *hars-1(RNAi)* (77%), *cars-1(RNAi)* (76%), and *kars-1(RNAi)* (67%) strains. Interestingly, the mitochondrial membrane potential showed more variable results between mt-ARS deficient strains when normalized to their reduced mitochondrial content, with significant reduction seen in normalized TMRE/MTG fluorescence in *hars-1(RNAi)* and *cars-1(RNAi)* worms. Interestingly, normalized TMRE/MTG fluorescence was significantly increased in several strains, with greatest increase as a metric of integrated OXPHOS capacity seen in *ears-2(RNAi)*, *dars-2(RNAi)*, and *yars-2(RNAi)* worms (Figure 2D). TMRE and MTG fluorescence measurements for all nineteen clones are presented in Figure S1A and S1B, respectively, and the corresponding TMRE data normalized to worm length (time of flight) is shown in Figure S1C.

Collectively, these data demonstrate that mt-ARS protein deficiencies induced by exposure to continuous feeding RNAi for one generation trend toward causing mitochondrial depletion, with more variable effect on mitochondrial OXPHOS as assessed by relative quantitation of in vivo membrane potential.

### 19 mt-ARS *C. elegans* knockdown strains generated by feeding RNAi for two generations manifest more pronounced disease phenotypes

As several mt-ARS knockdown strains surprisingly showed no obvious defect at the level of neuromuscular defects and mitochondrial stress induction following one generation of feeding RNAi exposure, we postulated that residual protein from the parent (‘maternal egg effect’) diminished the effective severity of RNAi knockdown. Therefore, *C. elegans* were exposed two consecutive generations to their respective feeding RNAi clone for each *mt-ARS* (second-generation strains denoted as *(RNAi)*^2^), with disease phenotype quantitative analysis then performed at L4+1 day of adulthood. Of note, 5 mt-ARS deficiency strains were excluded that showed sterility following the first generation of RNAi knockdown (*cars-1(RNAi)*, *fars-2(RNAi)*, *hars-1(RNAi)*, *kars-1(RNAi)*, *mars-2(RNAi)*). Indeed, consistently decreased worm linear growth was seen following two generations of RNAi knockdown for all but 4 mt-ARS strains (*aars-1(RNAi)*^2^*, tars-1(RNAi)*^2^, *yars-2(RNAi)*^2^, *gars-1(RNAi)*^2^) (Figure 3A). Comparison of worm length effects between the first- and second-generation of RNAi knockdown worms is shown in Table 2. Similarly, neuromuscular activity was reduced for 11 of the 15 second-generation (*RNA*i)^2^-induced mt-ARS knockdown strains, with a range of 26-66% mean reduction relative to wild-type worms (Figure 3B). A higher degree of UPR^mt^ stress induction was achieved in the second-generation RNAi exposed strains compared to the first-generation of RNAi knockdown worms, as exemplified by visual inspection of *wars-2(RNAi)*^2^ worms (Figure 3C). UPR^mt^ stress induction was quantified for all second-generation RNAi knockdown strains, with a range of 30-475% induction of UPR^mt^ seen in the different mt-ARS deficient strains relative to wild-type worms (Figure 3D). Thus, the second generation of RNAi knockdown worms consistently showed more pronounced disease phenotypes compared to the first generation. A summary of phenotypes observed for the first and second generation of RNAi knockdown for all 19 mt-ARS genes is shown in Table 2.

**Figure 3.**
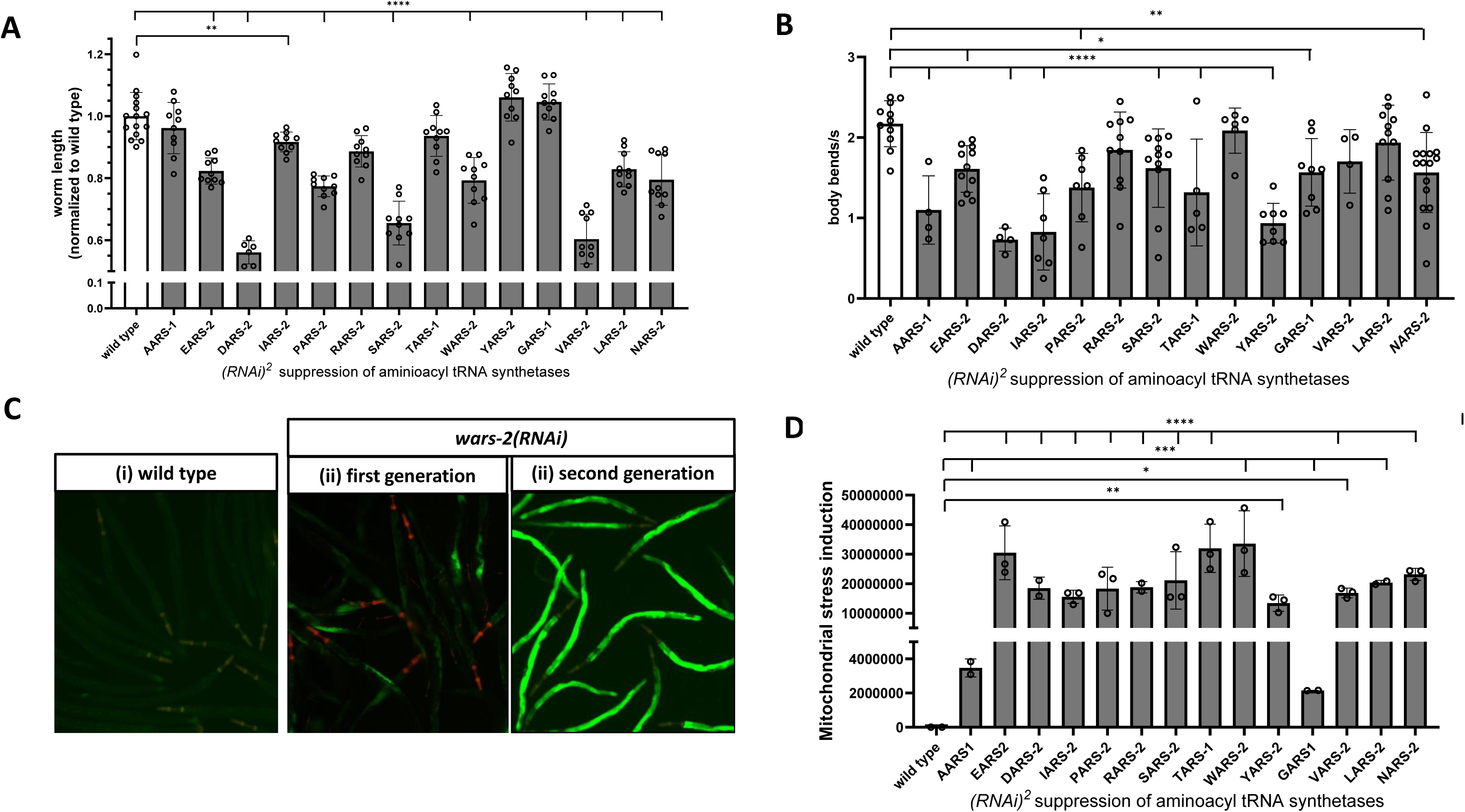
Phenotypic characterization of the second consecutive generation of mt-ARS RNAi knockdown worms. **(A)** Worm linear growth analysis shows reduced worm length in all knockdown strains relative to wild type. Data represent the mean ± SD of 10 individual animals. Statistical analysis by 1-way ANOVA with Dunnett post-hoc multiple comparisons. **(B)** Neuromuscular activity in the second generation of mt-ARS knockdown worms measured using a thrashing assay shows that the second generation has a stronger neuromuscular defect compared to the first generation (compare Fig 3B with 1B). Data represent the mean ± SD of 20 individual animals. Statistical analysis by 1-way ANOVA with Dunnett post-hoc multiple comparisons. **(C)** Representative images comparing the induction of UPR^mt^ stress in the first versus the second generation of *vars2 RNAi* knockdown. Shown are representative images of 3 biological replicates. **(D)** Quantification of UPR^mt^ stress in the second generation of RNAi knockdown, showing a higher degree of UPR^mt^ stress induction relative to the first generation of knockdown worms. Data represent the mean ± SD of 10 individual wells containing ∼50 worms. Variation was detected by 1-way ANOVA test. *: p <0.05, **: p<0.01, ***: p<0.001, ****: p<0.0001

### Cognate amino acid treatment rescued growth, neuromuscular activity, fertility, and UPR^mt^ stress inducion in mt-ARS knockdown worms

Recent studies of four ARS1-related cytosolic deficiencies demonstrated that human patient fibroblast models exhibit sensitivity to reduced levels of their respective cognate amino acids, with beneficial effect of high doses of oral cognate amino acid supplementation ^22,32^. However, it has been at matter of contention in the clinical community whether cognate amino acid supplementation is universally beneficial, without risk of adverse effects, in a ARS1 and ARS2 disorders. Therefore, we investigated the effects of cognate amino acid treatment in all 19 mt-ARS *C. elegans* RNAi knockdown models. Cognate amino acid treatment was continuously performed from embryo phase to L4+1 day stage of adulthood, using the viable and more severe of either the first- or second-generation of RNAi knockdown worms achieved for each gene (Figure 4A). Second-generation RNAi knockdown was used when there was no distinct thrashing or UPR^mt^ induction phenotype for first-generation RNAi worms (*gars-1(RNAi)*, *tars-2(RNAi)*, *nars-2(RNAi)*, *aars-1(RNAi)*, *kars-1(RNAi)* and *cars-1(RNAi)*).

**Figure 4.**
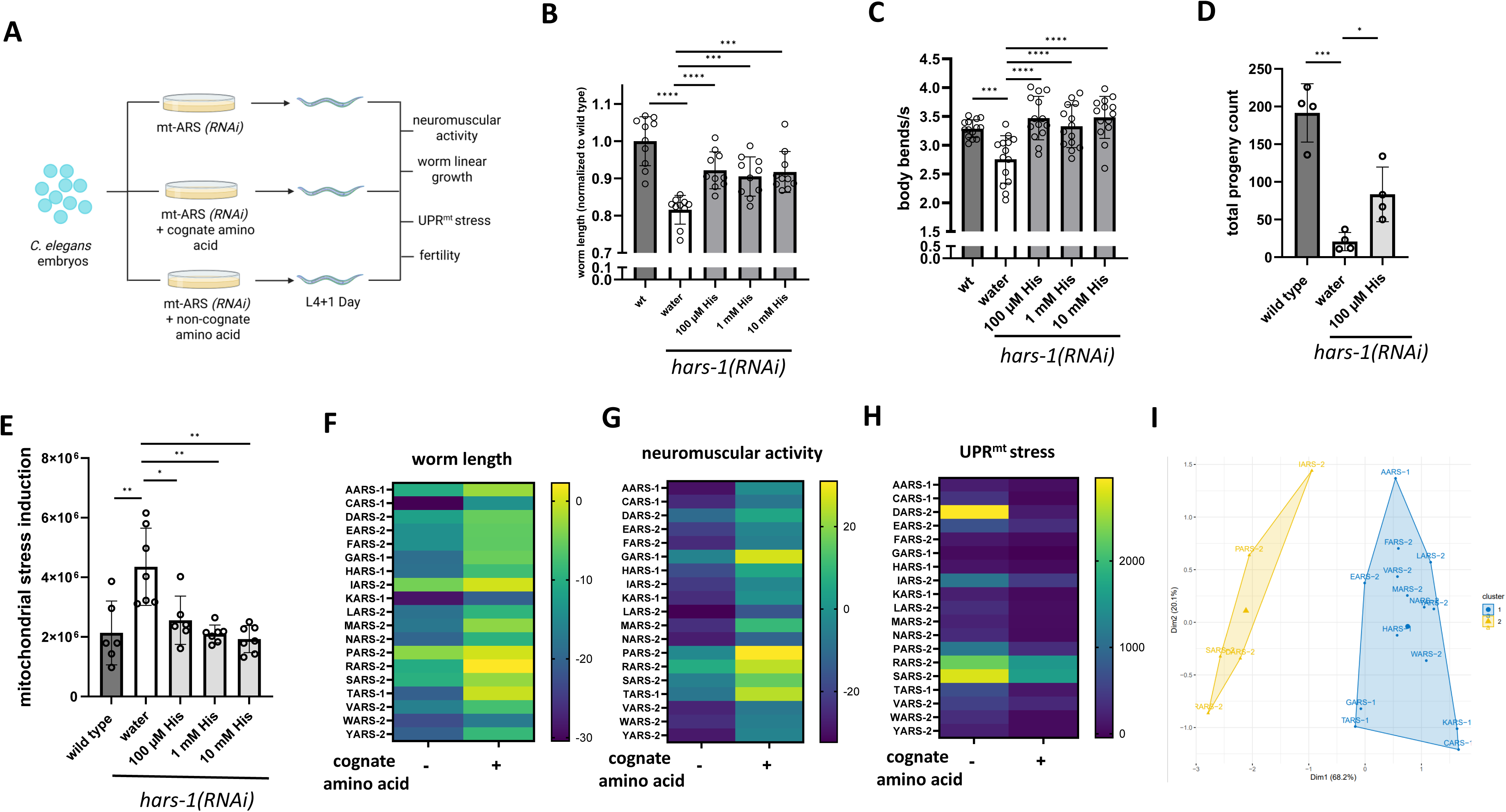
*hars-1* knockdown worm phenotypes are rescued in a dose-dependent fashion with their cognate amino acid, histidine. **(A)** Strategy for supplementation of knockdown worms with the cognate amino acid. Figure created with BioRender (https://www.biorender.com/) **(B)** Worm linear growth experiments show that treatment of *hars-1* knockdown worms with a 3-point dose curve (100 µM, 1 mM, 10 mM) of histidine leads to a significant increase in worm length at all tested concentrations. Data represents the mean ± SD of 10 individual animals. Variation was detected by Unpaired t test with Welch’s correction. **(C)** Neuromuscular activity of **hars-1(RNAi)** knockdown worms following treatment with a three-point histidine dose curve (100 µM, 1 mM, 10 mM) showed a significant increase in activity at all tested concentrations. Data represent the mean ± SD of 10 individual animals. Statistical significance was assessed using an unpaired *t*-test with Welch’s correction. **(D)** Progeny count assay demonstrated a significant increase in number of progeny produced by the *hars-1* knockdown worms following treatment with 100 µM histidine. Data represent the mean ± SD of 4 biological replicates. Statistical significance was assessed using an unpaired *t*-test with Welch’s correction. **(E)** UPR^mt^ stress stress measurements indicate that treatment of *hars-1(RNAi)* worms with a three-point histidine dose curve (100 µM, 1 mM, 10 mM) significantly reduced mitochondrial stress at all tested concentrations. Data represents the mean ± SD of 10 individual wells containing ∼50 worms. **(F)** Effect of cognate amino acid treatment on worm length across all 19 mt-ARS genes. **(G)** Effect of amino acid treatment on neuromuscular activity across all 19 mt-ARS genes **(H)** Effect of amino acid treatment on UPR^mt^ stress across all 19 mt-ARS genes. *: p <0.05, **: p<0.01, ***: p<0.001, ****: p<0.0001

First, the effect of a 3-log scale dose curve (100 µM, 1 mM, 10 mM) involving histidine treatment was evaluated for the first generation of *hars-1(RNAi)* knockdown worms on animal length, neuromuscular thrashing activity, fertility, and UPR^mt^ stress response. Histidine treatment was administered for three days, from the embryonic stage to L4+1 of adulthood. All tested histidine concentrations led to a significant increase in worm body length, with an average linear growth enhancement of approximately 11% (Figure 4B, Figure S2A). In addition, histidine treatment significantly improved neuromuscular thrashing activity, restoring it to levels comparable to wild-type controls. (Figure 4C). Interestingly, treatment with 100 µM histidine significantly rescued the fertility defect phenotype of the *hars-1(RNAi)* knockdown worms, resulting in a 30% increase in progeny production compared to the untreated control (Figure 4D, Figure S2B). In addition, treatment of *hars-1(RNAi)* worms with all tested concentrations of histidine decreased UPR^mt^ stress, restoring it to the levels measured in the wild-type worms (Figure 4E).

The effect of treatment with 100 µM cognate amino acid was then studied in all 19 mt-ARS RNAi knockdown strains generated. As described above for the *hars-1(RNAi)* knockdown strain, treatment was performed from embryo phase, with outcome measurements performed at stage L4+1 (the first day of adulthood). Importantly, no gross morphological toxicity was observed after treatment with any cognate amino acid. However, treatment with 1 mM and 10 mM aspartate or 1 mM and 10 mM glutamic acid led to an increase in mitochondrial stress in the *dars-2(RNAi)* and *ears2(RNAi)* knockdown strains, respectively (Figure S2C, Figure S2D). Overall, treatment with cognate amino acids improved disease phenotypes across all 19 mt-ARS knockdown strains, as evidenced by reduced UPR^mt^ stress induction, enhanced neuromuscular (thrashing) behavior, increased worm length (growth), and, in the cases of *fars-2(RNAi)* and *hars-1(RNAi)* knockdown worms, improved fertility. In contrast, no improvement in the fertility defects was observed for *mars-2(RNAi)*, *kars-1(RNAi)* and *cars-1(RNAi)* knockdown strains. The relative degree of treatment effects of cognate amino acids on the 19 mt-ARS knockdown worms are comparatively shown in heatmaps on worm linear growth (Figure 4F), neuromuscular activity (Figure 4G), and UPR^mt^ stress induction (Figure 4H).

In attempt to group the mt-ARS knockdown strains according to shared phenotypic profiles, an unsupervised k-means clustering approach was applied to the baseline (untreated)dataset using worm length, UPR^mt^ stress, and neuromuscular activity as input variables. The optimal number of clusters was identified as two, based on both the Elbow and Silhouette methods (Figure S3A). The resulting clusters (Figure 4I) captured distinct phenotypic signatures among the mt-ARS genes. Cluster 1 comprised strains with knockdown of 14 genes encoding both mitochondrial and dual-localized tRNA synthetases. In these mt-ARS knockdown worms, phenotypic correlation between worm length and UPR^mt^ stress was weak (r=5.56×10^−4^), but moderately increased following cognate amino acid supplementation (r=0.3), suggesting a coordinated recovery as mitochondrial translation capacity was partially restored (Figure S3B, C) Cluster 2, which included RNAi knockdown worms of 5 genes with exclusive mitochondria-localized enzymes (*iars-2*, *pars-2*, *sars-2*, *rars-2* and *dars-2*), displayed a strong inverse correlation between worm length and UPR^mt^ stress (r=-0.96). Knockdown of these mt-ARS genes strongly induces mitochondrial stress response but, unexpectedly, only modestly affects worm growth. Following treatment with cognate amino acid supplementation, the correlation shifted toward a moderate positive value (r=0.32), comparable to the value observed in Cluster 1 strains after supplementation (Figure S3B, C). Together, these results show that while amino acid supplementation restores overt disease phenotypes across both mt-ARS clusters, the clusters differ in the correlation of worm length and mitochondrial stress response, which we postulate reflects distinct patterns of stress-growth regulation. Interestingly, direct correlations between either worm length or mitochondrial stress induction also improved post-treatment in both clusters 1 and 2 with neuromuscular function (Fig S3C, D).

### Aspartate cognate amino acid treatment restored mitochondrial OXPHOS levels in *dars-2(RNAi)* knockdown worms

Since the most severe disease phenotypes were observed in the first-generation *dars-2(RNAi)* knockdown worms, this model was used to investigate the status of integrated OXPHOS capacity upon treatment with its cognate amino acid, aspartate. To confirm DARS-2 protein knockdown efficiency, western immunoblot analysis was performed, which demonstrated 88% reduction in DARS-2 protein levels in worms fed *dars-2(RNAi)* for one-generation as compared to wild-type worms fed control *L4440(RNAi)* bacteria (Figure 5A, Figure 5B, Figure S4A). For the aspartate supplementation experiments, a negative treatment control was included, in which *dars-2(RNAi)* knockdown worms were treated with 100 µM alanine, an abundant non-cognate amino acid. Treatment was performed on solid media from the egg phase, as described above, and worms were analyzed at stage L4+1 day of adulthood. Similar to the observations in *hars-1(RNAi)* knockdown worms, treatment of *dars-2(RNAi)* knockdown worms with aspartate - but importantly not with alanine - resulted in a significant 7% average increase in worm length (Figure 5C, Figure S4B), restoration of neuromuscular activity to wild-type levels (Figure 5D), and a decrease of UPR^mt^ stress to baseline levels seen in wild-type worms (Figure 5E).

**Figure 5.**
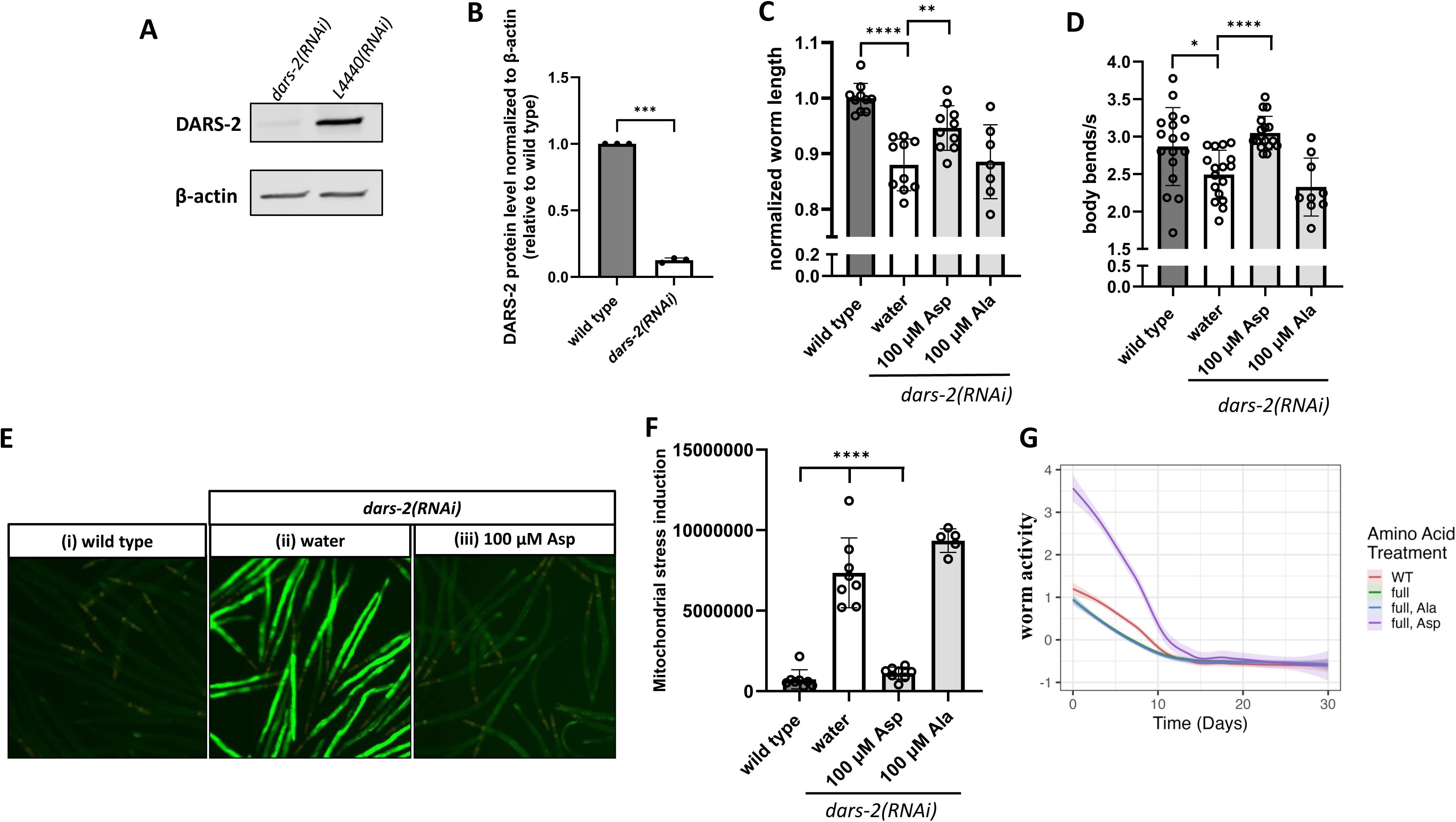
*dars2* knockdown worm phenotypes are rescued in a dose-dependent fashion by treatment with their cognate amino acid, aspartate. **(A)** Western Blot analysis of the level of DARS2 protein upon feeding *C. elegans* with *dars2 RNAi*. *L4440* bacteria was used as a negative control. A representative blot of three biological replicates is depicted. **(B)** Quantification of the Western Blot presented in **(A)**, confirming 87% level of DARS2 protein knockdown upon feeding *C. elegans* with *dars2 RNAi* bacteria. Data represents the mean ± SD of three individual biological replicates. Variation was detected by Unpaired t test with Welch’s correction. **(C)** Worm linear growth analysis of *dars2* knockdown worms shows a significant increase in worm length upon treatment with Asp compared to the water treatment as control. Data represents the mean ± SD of 10 individual animals. Variation was detected by Unpaired t test with Welch’s correction. **(D)** Neuromuscular activity of the *dars2* knockdown worms after treatment with 100 µM Asp (cognate aminoacid), 100 µM Ala (non-cognate aminoacid), or water (untreated control) shows a significant increase in thrashing activity after treatment with Asp. Data represents the mean ± SD of 20 individual animals. Variation was detected by Unpaired t test with Welch’s correction. **(E)** Representative high-content images of *dars2* knockdown worms treated with 100 µM Asp, 100 µM Ala or water show a decrease in GFP fluorescence upon treatment with Asp. **(F)** Quantification of UPR^mt^ stress of the dars2 knockdown worms after treatment with100 µM Asp, 100 µM Ala or water shows a significant rescue upon treatment with the cognate aminoacid (Asp), but not with the non-cognate aminoacid (Ala). Data represents the mean ± SD of 10 individual wells containing ∼50 worms. Variation was detected by Unpaired t test with Welch’s correction. **(G)** Worm activity measured using an automated lifespan imaging system revealed a significant increase in the activity of *dars-2(RNAi)* knockdown worms following treatment with 100 µM aspartate, whereas treatment with 100 µM alanine had no detectable effect. *: p <0.05, **: p<0.01, ***: p<0.001, ****: p<0.0001

Additionally, we evaluated both the median lifespan and healthspan of the *dars2(RNAi)* knockdown worms following supplementation with 100 µM aspartate or 100 µM alanine. To achieve this, we employed an automated platform designed to continuously monitor the movement of multiple *C. elegans* populations on solid media throughout their entire lifespan in a 24-well plate format (Figure S5A) ^33^. Treatment with 100 µM aspartate resulted in a noticeable increase in worm activity at days 2, 5, and 10 of adulthood (Figure 5G, purple curve) relative to untreated controls (Figure 5G, green curve) and wild type (Figure 5G, red curve). In contrast, supplementation with 100 µM alanine did not produce a similar effect (Figure 5G, blue curve). In addition, treatment with aspartate, but not alanine, resulted in a significant increase in the worm fraction (the proportion of the well area occupied by worms), consistent with the observed increase in worm size following aspartate supplementation (Figure S5D). Worm activity measurements for all treatment conditions are provided in Supplementary Fig. S4D. Despite the early improvement in activity observed with aspartate treatment at day 2 and 5 of adulthood (Figure S5C), no significant differences were detected in the median healthspan or lifespan of the *dars-2(RNAi)* worms (Figure S5D), indicating that the beneficial effects of aspartate on worm activity are limited to the early stages of adulthood rather than extending median survival and movement measured across 30 days.

Quantification of OXPHOS subunit levels by western immunoblot analysis was performed in the *dars-2(RNAi)* worms, both before and after treatment with aspartate or alanine. To detect the OXPHOS components, we used the total OXPHOS antibody (Abcam), which out of the five OXPHOS complexes, reliably quantifies *C. elegans* subunits of complex I, complex III, and complex V. TOM-20 is an outer mitochondrial membrane marker used as a protein control marker of mitochondrial content and β-actin intensity was used as loading control (Figure 6A, Figure S6). Worms fed *dars-2(RNAi)* bacteria and treated with water as buffer control showed a 50% increase in TOM-20 expression compared to worms fed *L4440* bacteria, indicating that knockdown of the DARS-2 protein triggers mitochondrial proliferation in *C. elegans*. Interestingly, worm treatment with 100 µM aspartate, but not with 100 µM alanine, rescued TOM-20 protein levels (Figure 6B). Interestingly, despite the increase in TOM-20 levels, Mitotracker Green fluorescence normalized to time-of-flight was reduced in the *dars-2(RNAi)* worms (Figure 2C), suggesting that mitochondria may have impaired membrane potential or altered morphology, which limits dye accumulation in the *dars-2* knockdown worms. Additionally, *dars-2(RNAi)* knockdown worms showed ∼50% reduced complex I subunit expression relative to wild-type worms, which was restored upon treatment with 100 µM aspartate but not 100 µM alanine (Figure 6C). Similarly, the protein levels of a complex III subunit were reduced to 50% in the *dars-2(RNAi)* knockdown worms, with restoration to wild-type levels upon treatment with 100 µM aspartate (Figure 6D). No significant change was observed complex V protein expression level in the *dars-2(RNAi)* knockdown worms (Figure 6E). Overall, these data confirm an outer mitochondrial membrane proliferation response at the level of TOM20 with reduction of OXPHOS subunits I and III expression, all of which are rescued by 100 µM aspartate cognate amino acid therapy in the first-generation *dars-2(RNAi)* knockdown worms.

**Figure 6.**
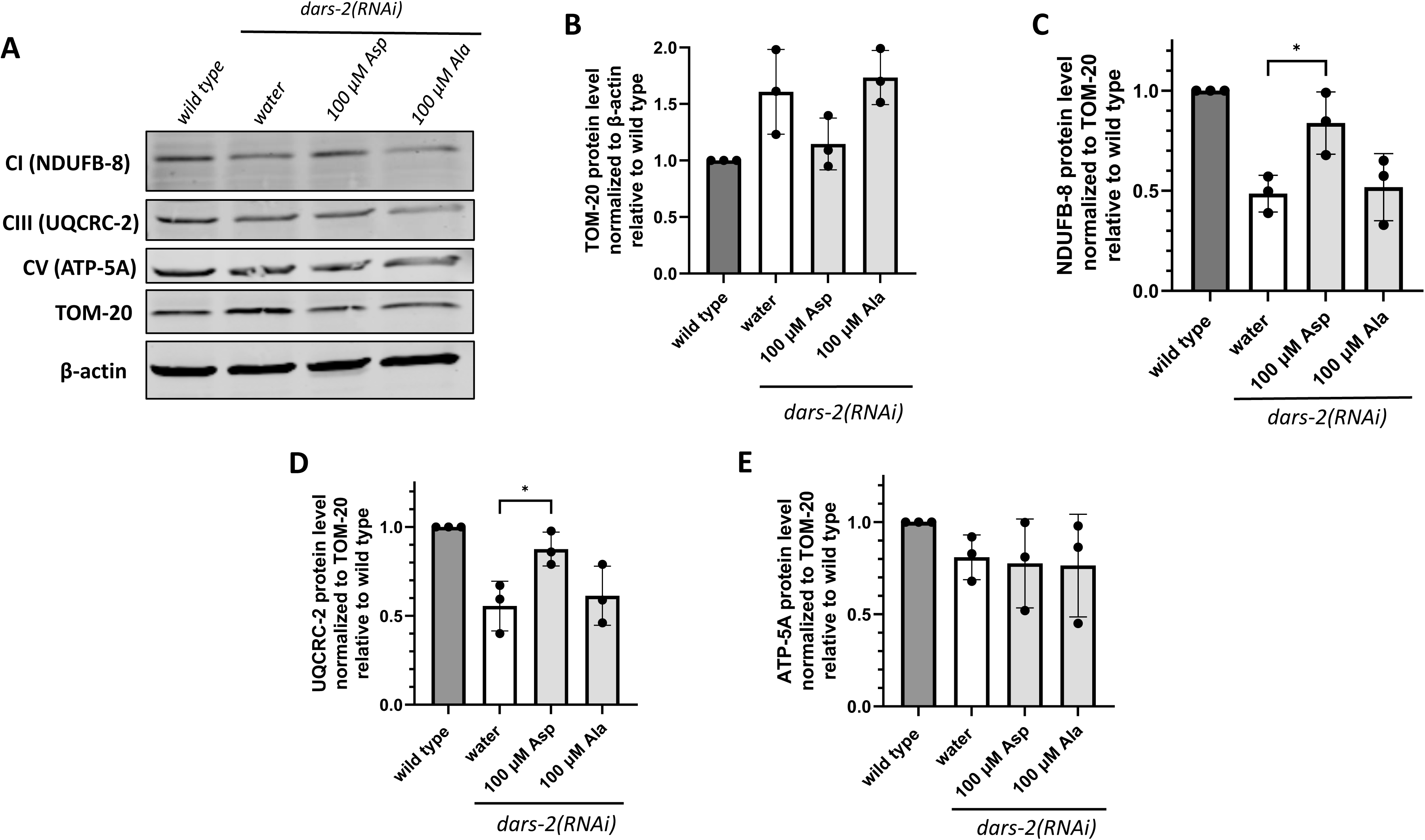
Mitochondrial OXPHOS subunit expression is specifically rescued in *dars2* knockdown worms by aspartate treatment. **(A)** Representative Western Blot images of mitochondrial proteins in the *dars2* knockdown worms after treatment with 100 µM Asp, 100 µM Ala and water. Shown are representative images of three individual biological replicates. **(B)** Quantification of Tom20 protein levels in the *dars2* knockdown worms treated with 100µM Asp or 100µM Ala. Tom20 protein levels were normalized to β-actin, and to wild type. Data represents the mean ± SD of three individual biological replicates. Variation was detected by Unpaired t test with Welch’s correction. **(C)** Quantification of the NDUFB8 protein as a marker of Complex I shows a decrease in Complex I upon *dars2* knockdown. Levels of Complex I were significantly increased after treatment with the cognate aminoacid (Asp) and were unchanged after treatment with the non-cognate aminoacid (Ala). Level of NDUFB8 protein were normalized to Tom20 and to the wild type control. Data represents the mean ± SD of three individual biological replicates. Variation was detected by Unpaired t test with Welch’s correction. **(D) (E)** Quantification of Complex III in the *dars2* knockdown model shows a significant decrease in UQCRC2 protein levels relative to wild type. Treatment with 100 µM Asp caused a significant increase in Complex III. No significant change in Complex II was observed after treatment with 100 µM Ala. Level of UQCRC2 protein were normalized to Tom20 and to wild type. Data represents the mean ± SD of three individual biological replicates. Variation was detected by Unpaired t test with Welch’s correction. **(F)** Quantification of Complex V shows no significant change in ATP5A levels upon *dars2* knockdown. Level ATP5A protein were normalized to Tom20 and to wild type. Variation was detected by Unpaired t test with Welch’s correction. *: p <0.05, **: p<0.01, ***: p<0.001, ****: p<0.0001

## DISCUSSION

To investigate the underlying phenotypic diversity and pathophysiology of mitochondrial tRNA synthetase disorders, we utilized feeding RNAi interference across one or two generations to systematically analyze deficiencies in all 19 genes that encode conserved mt-ARS proteins in *C. elegans*. Expression knockdown of each mt-ARS gene consistently resulted in linear growth deficiency, as well as decrease in mitochondrial mass in all worm strains except for AARS1. Interestingly, the greatest decrease in linear growth was observed upon knockdown of 3 *C. elegans* proteins having dual localization and function in both the mitochondria and cytoplasm: *hars-1*, *kars-1*, and *cars-1*. Across all 19 RNAi knockdown mt-ARS worm strains, variable effects were observed on lifespan, fertility, neuromuscular function (thrashing), UPR^mt^ stress induction, and relative mitochondrial membrane potential as a proxy for integrated OXPHOS capacity. For example, UPR^mt^ stress induction was most dramatically increased in DARS-2 deficient *C. elegans*. mt-ARS disease phenotypes were more pronounced in the second generation of RNAi knockdown worms relative to the first generation across all 19 mt-ARS knockdown strains, which likely reflects a maternal egg effect contributing greater residual ARS2 protein levels in the first as compared to second generation. Since the second consecutive generation of RNAi knockdown strains would minimize maternal egg effect, these findings suggest that the severity of disease phenotypes, particularly the activation of mitochondrial stress response and impairment of neuromuscular activity, correlates with the progressive decrease in residual levels of mt-ARS protein in *C. elegans*. Cognate amino acid therapies consistently and specifically rescued diverse phenotypes across each of the mt-ARS RNAi knockdown strains studied, including linear growth, neuromuscular activity, fecundity in some cases (*fars-2, hars-1*), and UPR^mt^ stress induction in the mt-ARS deficient worms. Maximal benefit was seen at 100 µM histidine cognate amino acid in a dose-ranging study of *hars-1* worms, with no adverse gross morphology-level effects observed. Cognate amino acid therapy further rescued mitochondrial OXPHOS complex I and III protein subunit expression in first-generation *dars-2(RNAi)* knockdown worms, and normalized elevated TOM20 levels that reflect their mitochondrial proliferation adaptive response. Collectively, these data demonstrate the utility of preclinical disease modeling by feeding RNAi knockdown in *C. elegans* across the entire class of mt-ARS disease. Study results further highlight that cognate amino acid supplementation is a promising precision therapeutic strategy across the class of mt-ARS deficiencies, which requires careful evaluation of their efficacy and safety in human clinical trials.

Although mt-ARS proteins are ubiquitously expressed and have similar functions in mitochondrial translation, disease syndromes and precise clinical manifestations caused by mutations in distinct mt-ARS genes are highly heterogeneous and tissue-specific. The underlying mechanisms responsible for this phenotypic variability and tissue specificity remain unclear, posing a significant challenge to the development of effective therapeutic strategies. Previous mouse experiments showed that loss of DARS2 triggers the activation of tissue-specific stress responses ^34^. In the heart, DARS2 ablation leads to the accumulation of unfolded and misfolded proteins, which activates strong adaptive stress response mechanisms independent of mitochondrial dysfunction. In contrast, no stress response activation was observed in skeletal muscle, suggesting that in mice, mitochondrial proteostasis and induction of mitochondrial stress are regulated in a tissue-dependent manner. Additionally, brain-specific DARS2 knockout in mice induced neuron-specific effects, leading to neuroinflammation and severe neurodegeneration in the forebrain and hippocampal neurons, while these effects were absent in oligodendrocytes despite their having significant mitochondrial impairment ^35,36^. Similarly, a mouse model carrying a hypomorphic point mutation in *WARS2*, *Wars2^V117L^*, exhibited tissue-specific alterations in mitochondrial biogenesis, steady-state OXPHOS levels, and heart-specific activation of stress response pathways ^17^. Collectively, these findings suggest that the tissue-specific phenotypes associated with mt-ARS-related diseases might arise at least in part from differential activation of stress response pathways and the varying capacities of distinct cell types to cope with impaired mitochondrial function. Similarly, our research group previously reported that tissue-specific activation of proteotoxic stress response occurs as a central mediator and therapeutic target of the pathophysiology underling variable disease phenotypes in a range of other genetic models of primary OXPHOS dysfunction, ranging from human cell line to *C. elegans* and mouse models^37,38^.

mt-ARS protein alterations caused by pathogenic mutations or reduced steady-state levels are likely to cause insufficient or mis-aminoacylation of mt-tRNAs ^39^, resulting in the synthesis of aberrant mitochondrial proteins and ultimately, to combined OXPHOS enzyme complex deficiencies. Unlike genetic knockouts that fully deplete mt-ARS protein expression, the feeding RNAi knockdown worm strains retain residual mt-ARS protein synthesis capacity. Therefore, we hypothesized that treatment of mt-ARS knockdown worms with their cognate amino acid would promote residual enzyme aminoacylation capacity, increasing the formation of correctly charged mitochondrial aminoacyl-tRNAs and restoring downstream mitochondrial OXPHOS function. Indeed, a study of four human fibroblast cell lines derived from ARS1 (cytosolic ARS deficiency) patients (*IARS1*, *LARS1*, *FARS1* and *SARS1*) demonstrated sensitivity to cognate amino acid deprivation, where oral treatment with the corresponding amino acids resulted in beneficial effects in all four ARS1 patients ^22^. Cognate amino acid supplementation similarly showed beneficial effects in human patients with pathogenic variants with *MARS1*^40–42^, in several humanized yeast models of *HARS1* deficiency^43,44^, and in one patient with *FARS2* deficiency ^45^. Moreover, beneficial effects have been recently described in 3 xenopus models of *VARS2* deficiency following treatment with valine ^46^. However, these studies describe individual ARS disorder and/or human patient treatment case reports, without any report of prior systematic investigation of amino acid supplementation on animal models across all mt-ARS deficiencies. Results of cognate amino acid therapy studies across a wide range of human patient primary fibroblast cell line *in vitro* models have been conflicting (Johan Van Hove, MD, PhD, personal communication). Furthermore, no natural history study has been performed across all mt-ARS disorders, nor have standardized clinical protocols or randomized, placebo-controlled clinical trials been reported of cognate amino acid therapies across mt-ARS disorders. The preclinical data identified here suggests future multi-site clinical trials of cognate amino acid therapies are warranted across the entire class of individually rare mt-ARS disorders, with careful consideration of the therapeutic impact of causal gene etiology, residual mt-ARS enzyme capacity, as well as complex aspects of clinical trial design to assess safety, tolerability, pharmacology, and efficacy on diverse multi-system phenotypes.

Having confirmed that the first- and second-generation of feeding RNAi knockdown worms were robust translational models that recapitulated major animal-level phenotypes and mitochondrial pathophysiologic aspects of human mt-ARS deficiencies, we systematically investigated the potential toxicity and therapeutic effect of cognate amino acids on objective disease phenotypes. Notably, treatment with 100 µM of each cognate amino acid in the 19 mt-ARS knockdown worm strains rescued diverse disease phenotypes, leading to increased animal linear growth and neuromuscular activity, improved animal fertility, and reduced mitochondrial stress. Moreover, treatment effects were shown to be specific to cognate amino acids. For example, the *dars-2(RNAi)* knockdown strain treatment only with aspartate, but not alanine, significantly rescued their reduced healthspan, increased mitochondrial mass, and restored OXPHOS subunit protein expression. Importantly, we did not observe any gross morphological toxicity after treatment with the cognate amino acid. However, very high concentrations of aspartate and glutamic acid were found to further increase mitochondrial UPR^mt^ stress induction in both wild-type worms and in *dars-2(RNAi)* knockdown strains. Further mechanistic studies are required to determine the underlying mitochondrial translation process components responsible for the beneficial effects observed following cognate amino acid supplementation in mt-ARS knockdown worms. Enhancing substrate availability through amino acid supplementation might promote the aminoacylation reaction, leading to an increase in the levels of charged mitochondrial tRNAs. In addition, protein folding and/or stability might be improved by increasing the substrate concentration, since substrate binding often stabilizes protein conformations. Additionally, in mt-ARS disorders, impaired charging of tRNA with its cognate amino acid can lead to altered levels of the corresponding amino acid ^47^, which may be restored through supplementation. In all cases, impaired mt-ARS function in mitochondrial translation of the 13 mtDNA-encoded mRNAs encoding subunits of complexes I, III, IV, and V results in reduced mitochondrial OXPHOS capacity, which is clearly deficient in the mt-ARS diseases and was found to be corrected by cognate amino acid treatment at the levels of OXPHOS protein expression and integrated mitochondrial membrane potential that relies upon intact OXPHOS activity.

Phenotypic analysis across all RNAi worm strains demonstrated that mt-ARS gene knockdown strains separates into two clusters, at baseline, with discrepantcorrelation patterns between mitochondrial stress induction and worm growth. Cluster 1, containing strains that inhibited both mitochondrial tRNA specific and dual-localized mitochondrial tRNA synthetase enzymes, showed very weak correlations between growth and mitochondrial stress induction, which increased modestly after cognate amino acid supplementation to suggest partial restoration of coordinated cellular function. In contrast, Cluster 2, composed entirely of worm strains with knockdown of mitochondria-localized enzymes, at baseline showed a strong inverse relationship between worm growth and mitochondrial stress, but upon amino acid supplementation, converges to yield the same moderate positive correlation between these variables as was seen post-treatment in Cluster 1. These findings suggest that the regulation of mitochondrial stress induction and linear animal growth is modulated differently across the two clusters. Nonetheless, cognate amino acid supplementation normalized the correlative relationship in both, as well their direct correlation with improved neuromuscular function. Notably, the negligible correlations at baseline (without treatment) between mitochondrial stress and worm length, as well as between mitochondrial stress and neuromuscular activity in Cluster 1 are suggestive that in addition to their roles in mitochondrial translation that these mt-ARS proteins may have moonlighting functions. Indeed, non-canonical functions have been attributed to AARS2^48,49^, WARS2^16^, TARS2^50^, all of which are members of Cluster 1. Additionally, within Cluster 1, knockdown of the four dual-localized (to support cytosolic and mitochondrial translation) ARS genes (*gars-1*, *tars-1*, *kars-1*, *cars-1*) resulted in the greatest decrease in worm length, while causing less mitochondrial stress induction and neuromuscular impairment than other genes in the cluster. This suggests that reduced cytoplasmic ARS expression may further limit animal growth due to insufficient cytoplasmic protein synthesis to support organismal growth.

In summary, our study provides the first systematic characterization of mt-ARS-associated disease phenotypes using *C. elegans* as a preclinical model animal and investigating effects over one or two generations of feeding RNAi knockdown to yield increasing degrees of mt-ARS deficiency. These translational models recapitulate key aspects of human disease phenotype, including impaired linear growth, decreased neuromuscular activity and progeny, and increased mitochondrial UPR^mt^ stress induction with reduced mitochondrial OXPHOS protein expression. Importantly, these data provide confirmatory evidence supporting the safety and efficacy of cognate amino acid 100 µM supplementation as a promising therapeutic strategy for all mt-ARS deficiencies, providing compelling rationale to pursue rigorous clinical trials in a targeted treatment approach for human mt-ARS disorders.

## Acknowledgements

This work was funded in part by the CureARS Foundation and the National Institutes of Health (NIH, R35-GM134863, M.J.F.). The content is solely the responsibility of the authors and does not necessarily represent the official views of the NIH.

## Author Contributions

MJF and NDM conceived of and designed the study and obtained study funding. MJF reviewed all study data and figures and assisted in manuscript preparation. CR performed all phenotypic characterization and amino acid treatment experiments. NM generated the double-transgenic hsp-6p::gfp + myo-2p::mCherry worm strain, performed WormScan lifespan analyses, trained personnel on high throughput methodology, and assisted with worm homology determination and phenotypic characterization. KK performed the automated lifespan data analyses. DMI performed all correlation analyses. EO contributed to experimental design, analysis, troubleshooting, and data interpretation. RX reviewed and advised on the statistical methods. CR, VEA, and MJF drafted the manuscript. All authors reviewed and approved the final manuscript.

## METHODS

### Reagents and tools

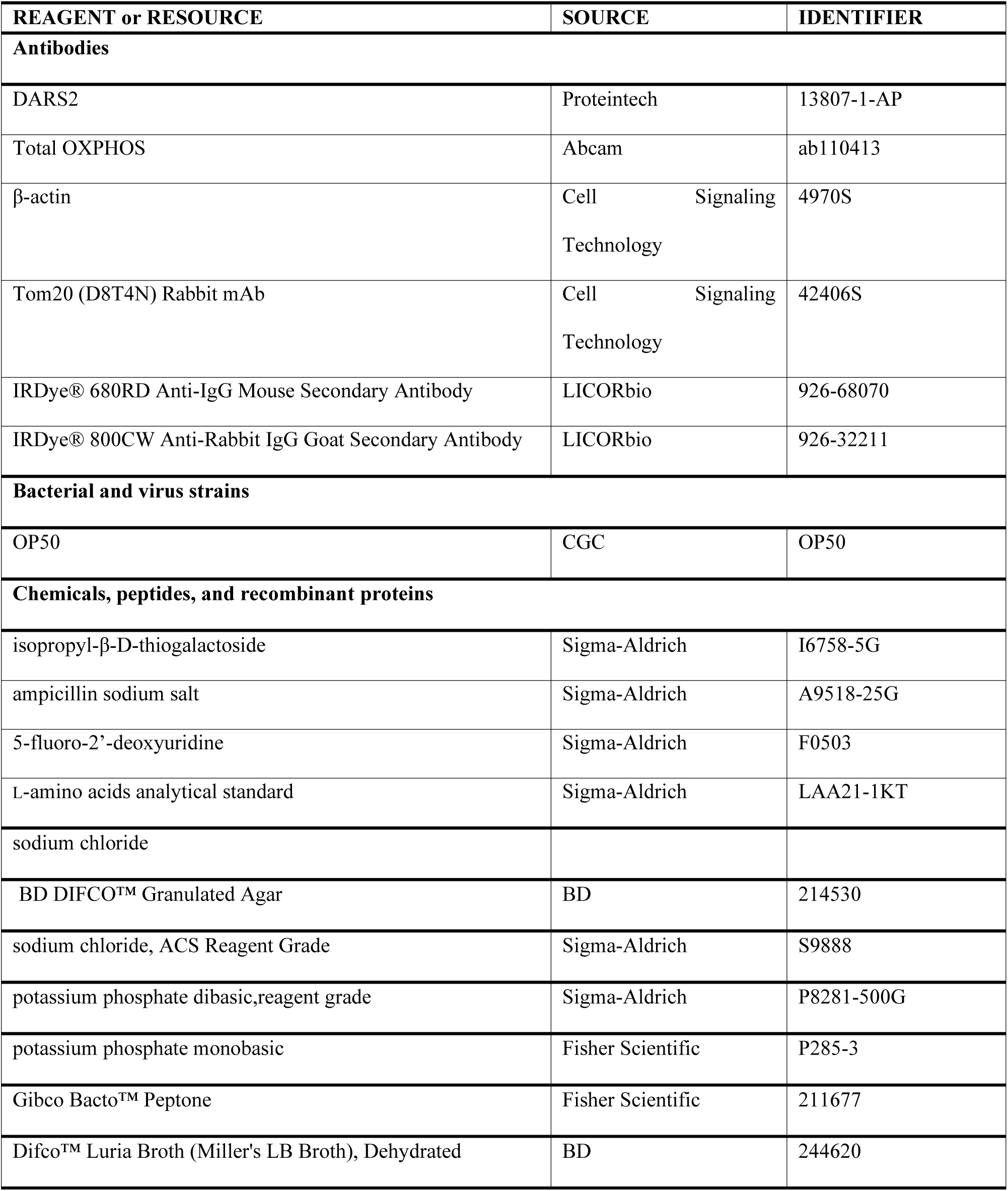
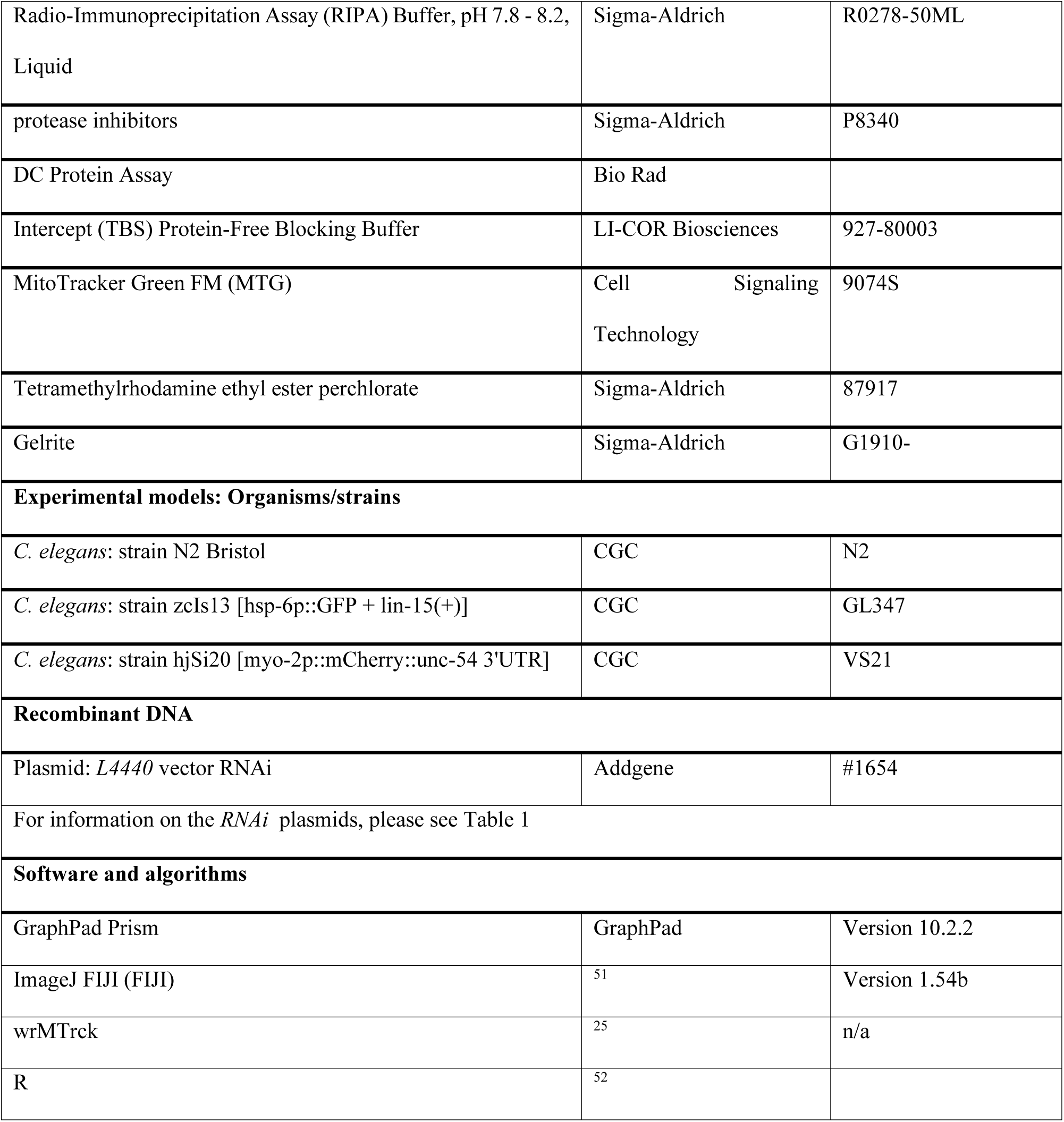

#### *C. elegans* growth and maintenance

The worm strain (GL347) with hsp-6p::gfp and worm strain (VS21) with (myo-2p::mCherry) was purchased from the Caenorhabditis Genetics Center (CGC) at the University of Minnesota. The strain was crossed in order to create a wild-type strain with (hsp-6p::gfp + myo-2p::mCherry). Worms were maintained at 20°C, on nematode growth media (NGM) agar plates seeded with OP50 bacteria.

#### Synchronization of *C. elegans*

Synchronous worm populations were obtained by bleaching. Worms were collected from NGM agar plates in 15 mL conical tubes with 4 mL S. Basal (S basal medium (23.4 g NaCl, 4 g K_2_HPO_4_, 25 g KH_2_PO_4_, 4 ml cholesterol (5 mg/ml in ethanol), 4 L milliQ water, pH = 7). Subsequently, 1 mL 100% commercial germicidal bleach (Clorox) and 500 µL 1 M NaOH were added and the tubes were gently mixed for approximately 2 min until no worm body was visible. Samples were centrifuged at 182*g* for 1 min, and the supernatant was immediately removed. The pellet was washed twice by resuspension in 10 mL S. Basal and centrifugation at 182*g* for 1 min.

#### Knockdown of mtARS genes in *C. elegans*

*RNAi* NGM agar plates were supplemented with 2 mM isopropyl β-D-1-thiogalactopyranoside (IPTG) and 100 µg/mL ampicillin. *RNAi* bacteria were inoculated in 30 mL Luria Broth (LB) containing 100 µg/mL ampicillin at 37°C overnight, concentrated 10 times and 500 µL resulting bacteria culture was seeded on *RNAi* NGM agar plates.

#### Treatment of mtARS knockdown *C. elegans* with the cognate amino acid

Worms were treated with the specified concentration of cognate amino acid on 100 mm *RNAi* NGM plates, from embryo phase. *RNAi* plates were prepared as described above, the desired concentration of amino acid was subsequently supplemented, and the plates were dried before seeding them with *RNAi* bacteria.

#### Locomotor behavior and activity analyses in *C. elegans*

Worm activity in liquid media was quantified using a thrashing assay. Worms were placed in 1 mL room temperature S. Basal and a 10 s video was recorded using a Basler USB camera (Basler AG, Germany, #108014) with a KOWA industrial lens 75mm/F2.5. The activity of the worms was analyzed in FIJI^51^, using a previously developed plugin ^25^. A total of three replicates with 20 animals per condition were performed.

#### Western Blotting

Worms (∼1000 animals) were grown on 150 mm NGM agar plates until stage L4+1 of adulthood and collected with 10 mL S. Basal. Samples were then centrifuged at 182*g* for 1 min and the supernatant was immediately removed. The resulting pellets were washed twice with 10 mL S. Basal and stored at -80°C. Worms were lysed by addition of an equal volume of RIPA buffer (Sigma-Aldrich) supplemented with 1 × protease inhibitors (Sigma-Aldrich) and homogenized on ice for 2 min using a pellet pestle. Samples were then centrifuged at 21300*g* for 20 min and the pellet was discarded. The total protein concentration of the supernatant was measured using a DC Protein Assay Kit (Bio-Rad) kit, following the manufacturer’s instructions. A total of 50 µg total protein/lane were separated on a Mini-PROTEAN® TGX™ Precast Protein Gels, 4–15%, 10-well using NuPAGE 1× MES (Thermo Fisher) running buffer. Proteins were then transferred onto nitrocellulose membranes (Bio-Rad) using 1× wet western blot transfer buffer (40 mM glycine, 50 mM Tris and 20% methanol) for 1 h at 350 mA at 4 °C. The membranes were blocked in 1 x Blocking Buffer (Intercept (TBS) Protein-Free Blocking Buffer, LICOR) for 1 h at room temperature and incubated with primary antibodies diluted in blocking buffer overnight at 4°C. Blots were washed with TBS with 0.2% Tween (TBS-T) three times for 10 min and further incubated with secondary antibodies for 1 h at room temperature. Membranes were washed three times with TBS-T for 10 minutes at room temperature and imaged using an Odyssey CLx Infrared Imaging System. The intensity of the bands was quantified using FIJI^51^.

#### Worm mitochondrial unfolded protein (UPR^mt^) stress assay using high content imaging (CX5)

For mitochondrial stress assays, we used the *C. elegans* strain expressing the hsp-6p::GFP and myo-2p::mCherry reporters. Mitochondrial stress was assessed by quantifying the induction of GFP fluorescence controlled by the *hsp-6p*::GFP reporter for the *C. elegans* homologue to human HSP70. Synchronized stage L4+1 Day worms were collected from NGM plates with S. Basal and the worm density was adjusted to 1 worm/µL. Fifty µL samples were dispensed into 384 well plates, to obtain a total of 50 worms/well. The CX5 SpotDetector was used to quantify the number of worm heads in the red channel, and the total green fluorescence in each well was used as an indicator of mitochondrial stress response induction. The total green fluorescence in each well was divided by the number of worms to quantify the mitochondrial stress response per worm.

#### Worm brood size assay

Worms were synchronized by bleaching. Stage L4 worms were transferred on 3.5 cm NGM plates (1 worm per plate) supplemented with 2 mM IPTG and 100 µg/mL ampicillin and seeded with the corresponding RNAi bacteria. For three consecutive days the worm was transferred to a fresh 3.5 cm plate. The number of progenies was counted on each plate the day after the worm had been transferred onto a fresh plate. At least three biological replicates per condition were performed.

#### Worm length analysis

*C. elegans* were imaged on NGM agar plates using a Basler USB camera (Basler AG, Germany, #108014) with a KOWA industrial lens 75mm/F2.5, and the length of each individual worm was analyzed using FIJI^51^. Three biological replicates of 10 animals were analyzed. The average length of the knockdown worms was normalized to the average length of the wild type worms.

#### Worm mitochondrial mass and membrane potential relative quantitation

Worms were co-labeled with tetramethylrhodamine ethyl ester (TMRE) as an indicator of mitochondrial membrane potential and MitoTracker Green FM (MTG) as an indicator of mitochondrial mass. Synchronized stage L4 worms were transferred to NGM agar plates containing 100 μM TMRE and 2 μM MTG for 24 h at 20°C. The worms were then transferred to NGM agar plates without fluorophores for 1 h at 20°C to allow the intestinal clearing of the fluorescent dyes. The fluorescent intensities of the red and green channels were then quantified using a Biosorter (Union Biometrica). The red fluorescence (membrane potential) was normalized to the green fluorescence (mitochondrial mass). Experiments were performed in triplicates, with 300 worms per condition.

#### Lifespan analyses by WormScan analsis

Worm lifespan was measured using the WormScan method ^26^. Animals were cultured on NGM agar plates supplemented with 100 μg/ml FUDR to a density of ∼50 worms/3.5 cm plate. Two consecutive images were acquired using an Epson V800 scanner every two days. The worms that moved after the first scan were identified and counted using FIJI^51^.

#### Automated lifespan and healthspan measurements by WormRobot analysis

Preparation of 24-well plates for automated lifespan measurements was performed as previously described^33^, with minor modifications. The media in between wells contained of 0.3% NaCl and 0.66% gelrite, while the medium in the wells contained 0.3% NaCl, 0.25% peptone, 0.8% gelrite, 50 µM FUDR, 2 mM IPTG, and 100 µg/mL ampicillin. The region between wells was filled with approximately 30 mL of in between wells media, and a total volume of 1 mL medium was dispensed into each well using a 10 mL automatic pipette. After the media solidified, wells were supplemented with either 100 µM aspartate, 100 µM alanine, or water as a control. To minimize edge effects and prevent non-uniform imaging artifacts, treatment conditions were randomized across the 24-well plate (Figure S5A). Each plate included six technical replicates for the following conditions: wild-type, *dars2(RNAi)* water control, *dars-2(RNAi)* + aspartate, and *dars-2(RNAi)* + alanine. Wells were subsequently seeded with the corresponding RNAi bacteria: *dars-2(RNAi)* or *L4440* control bacteria and dried overnight. Synchronized L4-stage worms, fed with *dars-2(RNAi)* or *L4440 (RNAi)* bacteria and treated from the embryonic stage with 100 µM aspartate, 100 µM alanine, or water (control), were prepared as described above. Approximately 30 worms were then transferred into each well. After allowing the wells to dry, the plate lids were treated with antifog solution and the plates were sealed with parafilm to prevent evaporation during the experiment. Two images of each plate were automatically captured twice daily—once before and once after blue light stimulation—as previously described^33^. Data was analyzed and visualized using R^52,53^. For activity, data is normalized to the worm fraction, then z-scored by condition. The day of max activity is found by taking the day where activity reached its absolute maximum for each animal. Healthspan and lifespan are calculated by rescaling all data from 0 to 1, then calculating total cumulative activity. Animals were considered to have reached their maximum healthspan at 82% of cumulative activity and their complete lifespan at 99% activity, as described in Fouad, et al 2021, who developed the machine and the image analysis software^53^. Comparisons between groups were analyzed using ANOVA with a post-hoc Tukey test. An adjusted probability (P) value of less than 0.05 was considered statistically significant. Plots were made using ggplot2^52^.

#### Quantification and statistical analysis

FIJI was used to perform the worm length, thrashing and lifespan analyses. For lifespan assays, survival graphs, i.e., Kaplan-Meier curves, were generated in GraphPad Prism. Log-rank test was used to compare the survival curves between conditions.

Except for the automated lifespan experiments, all other statistical analyses were performed in GraphPad Prism software, using t-tests or ANOVA with Brown-Forsythe and Welch tests.

Data are presented as means ± SD unless otherwise noted. Statistical parameters are reported in figures and corresponding figure legends.

To assess whether the genes are clustered by the observed phenotypes, an unsupervised learning approach, the k-mean clustering method implemented in R *kmeans* function, is applied to untreated worm data including the worm length, UPRmt stress, and neuromuscular activity. As the phenotypes were obtained from distinct worms, the mean of each phenotype was computed and used as input for the k-means clustering. The optimal number of clusters was determined by the Elbow graph and the silhouette method (Figure S3B).

Phenotypic relationships among mt-ARS knockdown worms were assessed using Pearson correlation coefficients. Correlations were calculated between measured phenotypes (worm length, neuromuscular activity, and mitochondrial stress) both before and after cognate amino acid supplementation. Analyses were performed for the full set of knockdowns as well as within each identified cluster. Statistical analyses and correlation computations were performed in GraphPad Prism.

The automated lifespan and healthspan experiments were analyzed and data was visualized using R. Details of the data processing and analysis are provided in the Methods section.

## Notes

**Conflict of Interest Statement**. MJF is inventor of US patent 12,011,452 B2 issued Jun 18, 2024, “Compositions and Methods for Treatment of Mitochondrial Respiratory Chain Dysfunction and Other Mitochondrial Disorders”. MJF is engaged with companies involved in mitochondrial disease therapeutic preclinical and/or clinical-stage development. MJF has served as co-founder and chief scientific advisor of Rarefy Therapeutics; an advisory board member with equity interest in RiboNova Inc.; a scientific advisory board member and paid consultant with Khondrion and Larimar Therapeutics; paid consultant for Ajinomoto Cambrooke, Inc, Astellas (formerly Mitobridge), Casma Therapeutics, Cyclerion Therapeutics, Epirium Bio (formerly Cardero Therapeutics), HealthCap VII Advisor AB, HTH Capital, Imel Therapeutics, Mayflower, Inc., Primera Therapeutics, Inc., Minovia Therapeutics, Mission Therapeutics, NeuroVive Pharmaceutical AB, Precision Biosciences, Reneo Therapeutics, Saol Therapeutics, Stealth BioTherapeutics, Vincere Bio, and Zogenix; and/or has been a sponsored research collaborator for Aadi Bioscience, Adjuvia Therapeutics, Astellas, Cyclerion Therapeutics, Epirium Bio, Imel Therapeutics, Khondrion, Merck, Minovia Therapeutics, Mission Therapeutics, NeuroVive Pharmaceutical AB, Precision Biosciences, PTC Therapeutics, Raptor Pharmaceuticals, REATA Inc., Reneo Therapeutics, RiboNova, Saol Therapeutics, Standigm, Stealth BioTherapeutics, and Thiogenesis. MJF also has received royalties from Elsevier. None of the other authors have relevant conflicts of interest to declare.

### Competing Interest Statement

MJF is inventor of US patent 12,011,452 B2 issued Jun 18, 2024, 'Compositions and Methods for Treatment of Mitochondrial Respiratory Chain Dysfunction and Other Mitochondrial Disorders'. MJF is engaged with companies involved in mitochondrial disease therapeutic preclinical and/or clinical-stage development. MJF has served as co-founder and chief scientific advisor of Rarefy Therapeutics LLC and M-Vortex LLC; an advisory board member with equity interest in RiboNova Inc.; a scientific advisory board member and paid consultant with Khondrion and Larimar Therapeutics; paid consultant for Ajinomoto Cambrooke, Inc, Astellas (formerly Mitobridge), BPGbio (with equity), Casma Therapeutics, Cyclerion Therapeutics, Epirium Bio (formerly Cardero Therapeutics), HealthCap VII Advisor AB, HTH Capital, Imel Therapeutics, Mayflower, Inc., Primera Therapeutics, Inc., Minovia Therapeutics, Mission Therapeutics, NeuroVive Pharmaceutical AB, Precision Biosciences, Reneo Therapeutics, Saol Therapeutics, Stealth BioTherapeutics, Vincere Bio, and Zogenix; and/or has been a sponsored research collaborator for Aadi Bioscience, Adjuvia Therapeutics, Astellas, Cyclerion Therapeutics, Epirium Bio, Imel Therapeutics, Khondrion, Merck, Minovia Therapeutics, Mission Therapeutics, NeuroVive Pharmaceutical AB, Precision Biosciences, PTC Therapeutics, Raptor Pharmaceuticals, REATA Inc., Reneo Therapeutics, RiboNova, Saol Therapeutics, Standigm, Stealth BioTherapeutics, and Thiogenesis. MJF also has received royalties from Elsevier and Chemistry Rx. None of the other authors have relevant conflicts of interest to declare.

