## supplemental files for "Cognate amino acid therapies provide preclinical benefit in 19 *C. elegans* models of ARS2 deficiency"

**Figure S1**

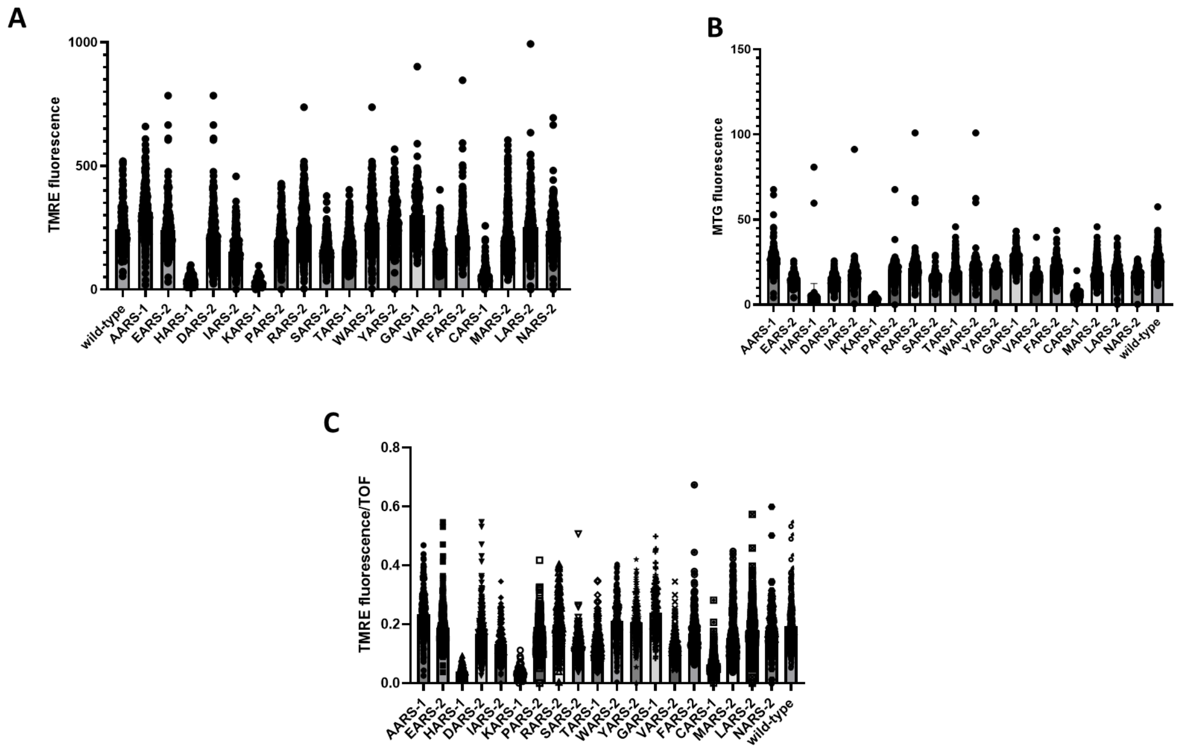


**Figure. S1. BioSorter analysis of mitochondrial function. (A)** TMRE fluorescence levels in the nineteen mt-ARS knockdown worm strains generated in the first generation of RNAi knockdown. Data represents the mean ± SD of 300 worms. **(B)** TMRE fluorescence levels in the nineteen mt-ARS knockdown worm strains generated in the first generation of RNAi knockdown. Data represents the mean ± SD of 300 worms. **(C)** TMRE fluorescence normalized to the time of flight in the nineteen mt-ARS knockdown worm strains generated in the first generation of RNAi knockdown. Data represents the mean ± SD of 300 worms. **Related to Fig. 2**

**Figure S2**

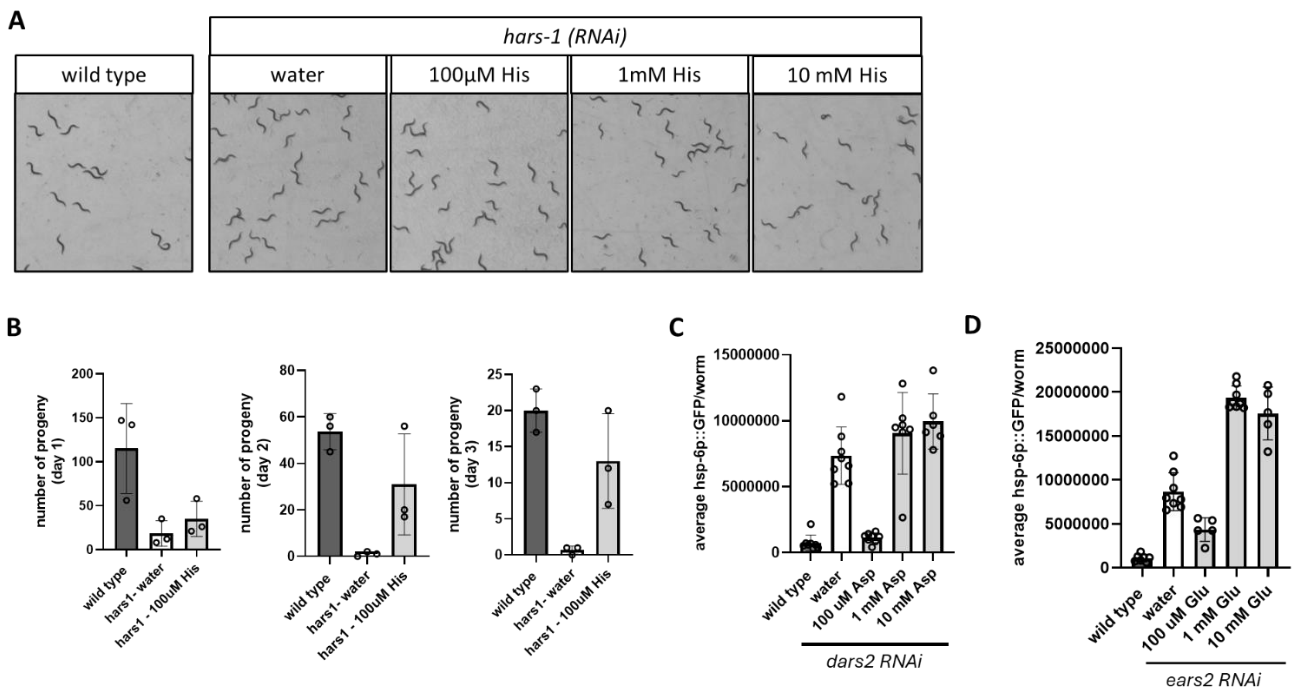


**Figure S2. Treatment of mt-ARS knockdown worms with their cognate amino acid. (A)** Representative images of *hars-1* knockdown worms treated with a 3-point dose curve of histidine (100 µM, 1 mM, 10 mM). L4440 RNAi worms were used as the wild type control, while treatment with water was used as the untreated control. **(B)** Number of progeny produced by *hars-1(RNAi)* knockdown worms treated with 100 µM histidine or water (untreated control), compared to wild-type worms. Worms were transferred to fresh plates, supplemented with the corresponding treatment daily throughout the experiment. **(C, D)** Treatment of *dars-2(RNAi)* and *ears-2(RNAi)* knockdown worms with a three-point dose curve of aspartate and glutamic acid, respectively, resulted in increased $UPR^{mt}$ stress at 1 mM and 10 mM concentrations. Data represent the mean ± SD of six wells, with approximately 50 worms per well. **Related to Fig. 4.**

Figure S3

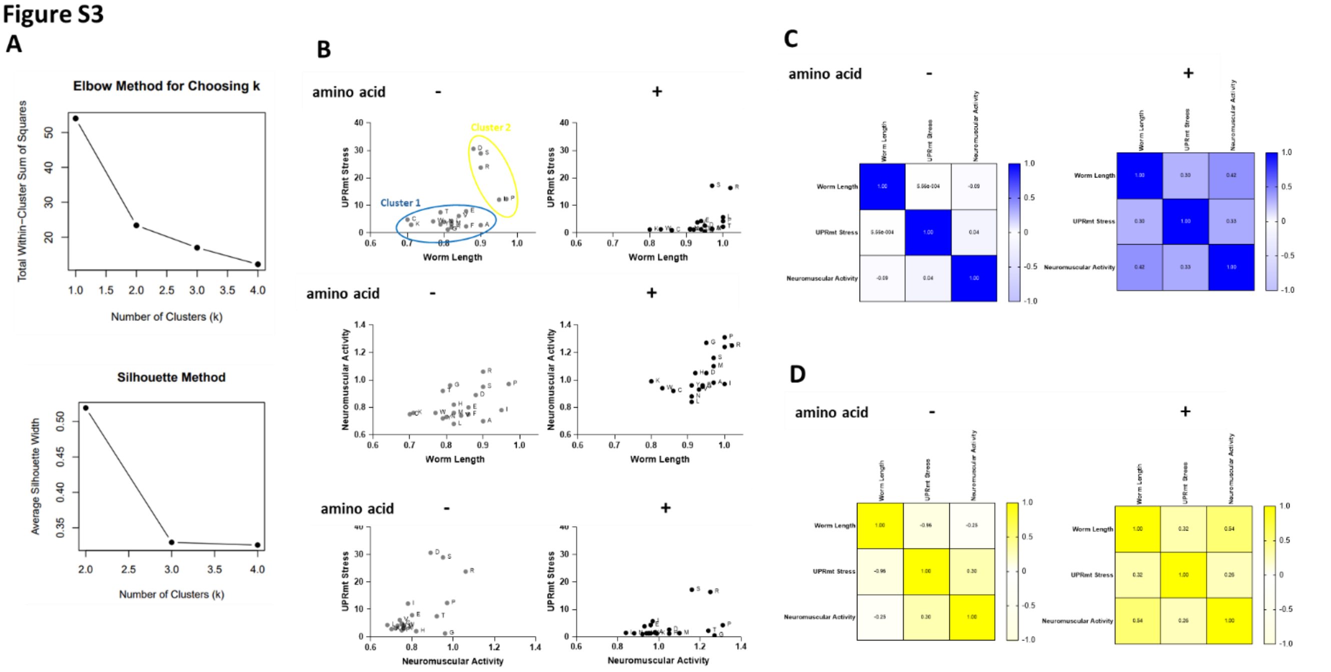


**Figure S3. Phenotypic separation of mt-ARS genes into two distinct clusters. (A)** Both the Elbow method and Silhouette method indicate that mt-ARS phenotypes can be optimally divided into two clusters. **(B)** Correlation analyses of measured phenotypes (worm length, neuromuscular activity, mitochondrial stress) for the full set of mt-ARS knockdowns, before and after cognate amino acid supplementation. **(C)** Within Cluster 1, phenotypic correlations are initially low, but show a moderate increase following cognate amino acid supplementation. **(D)** Within Cluster 2, mitochondrial stress and worm length exhibit a strong inverse correlation that shifts to a moderate positive correlation after amino acid supplementation, with correlation coefficients similar to the ones observed in Cluster 1. **Related to Figure 4.**

### Figure S4

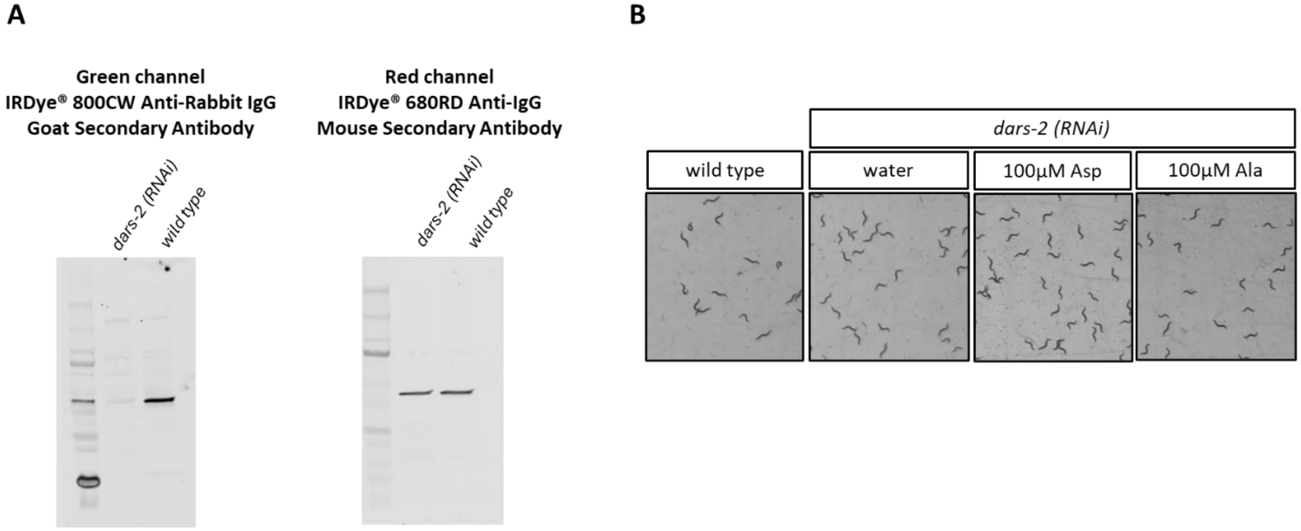


**Figure S4. Treatment of *dars-2* knockdown worms with aspartate. (A)** Full western immunoblot presented in Fig. 5A. **(B)** Representative images used for worm length analysis, of *dars-2* knockdown worms treated with 100 µM aspartate, 100 µM alanine or water (non-treated control) compared to wild type worms. **Related to Fig 5.**

**Figure S5**

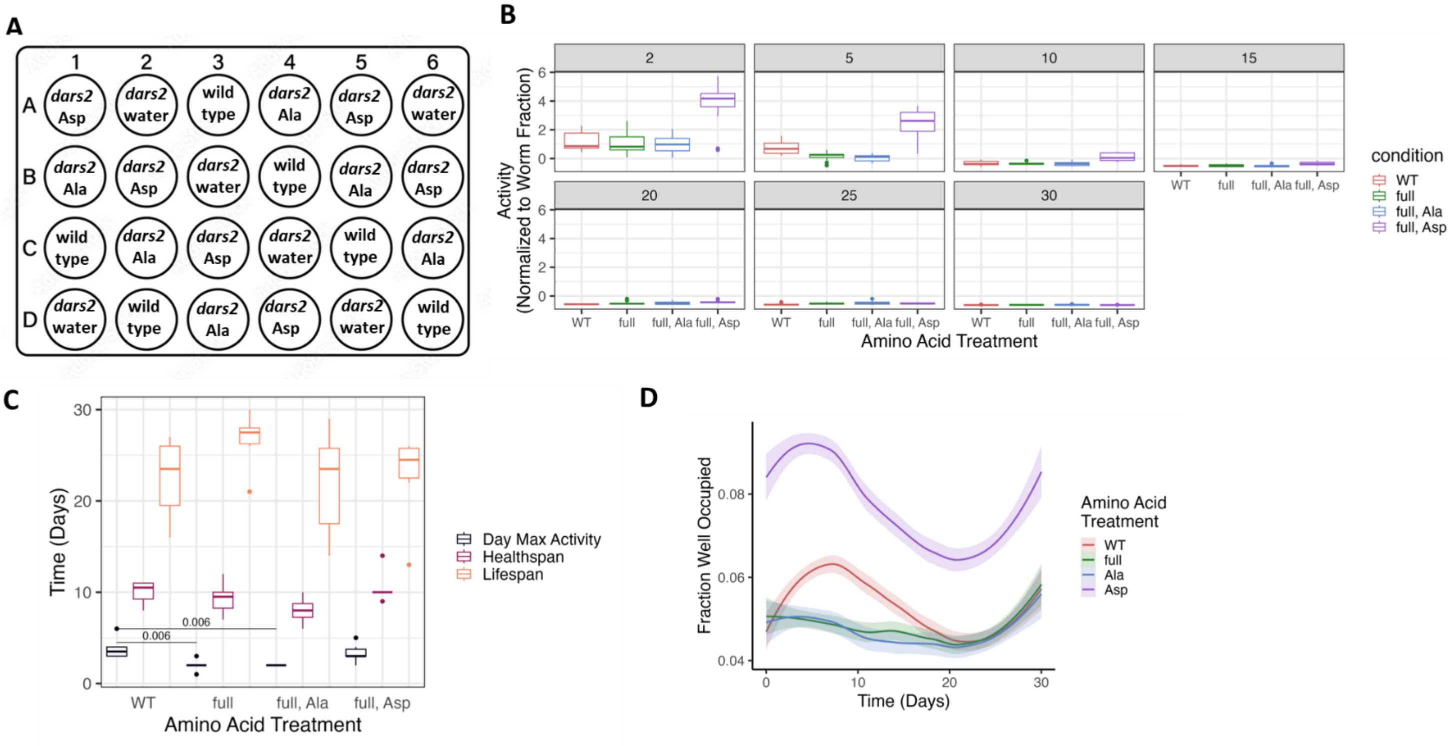


**Figure S5. Automated worm lifespan analyses on WormRobot. (A)** Plate map showing wild-type and *dars-2(RNAi)* knockdown worms treated with aspartate, alanine, or water. Test conditions were randomized across the 24-well plate to minimize edge effects. **(B)** Worm activity normalized to worm fraction shown for days 2, 5, 10, 15, 20, 25 and 30. **(C)** Mean healthspan, lifespan and day of maximum activity for the populations of worms described in **(A)**. **(D)** Worm fraction, which correlates with the size of the worms measured across 30 days. **Related to Fig. 5G.**

**Figure S6**

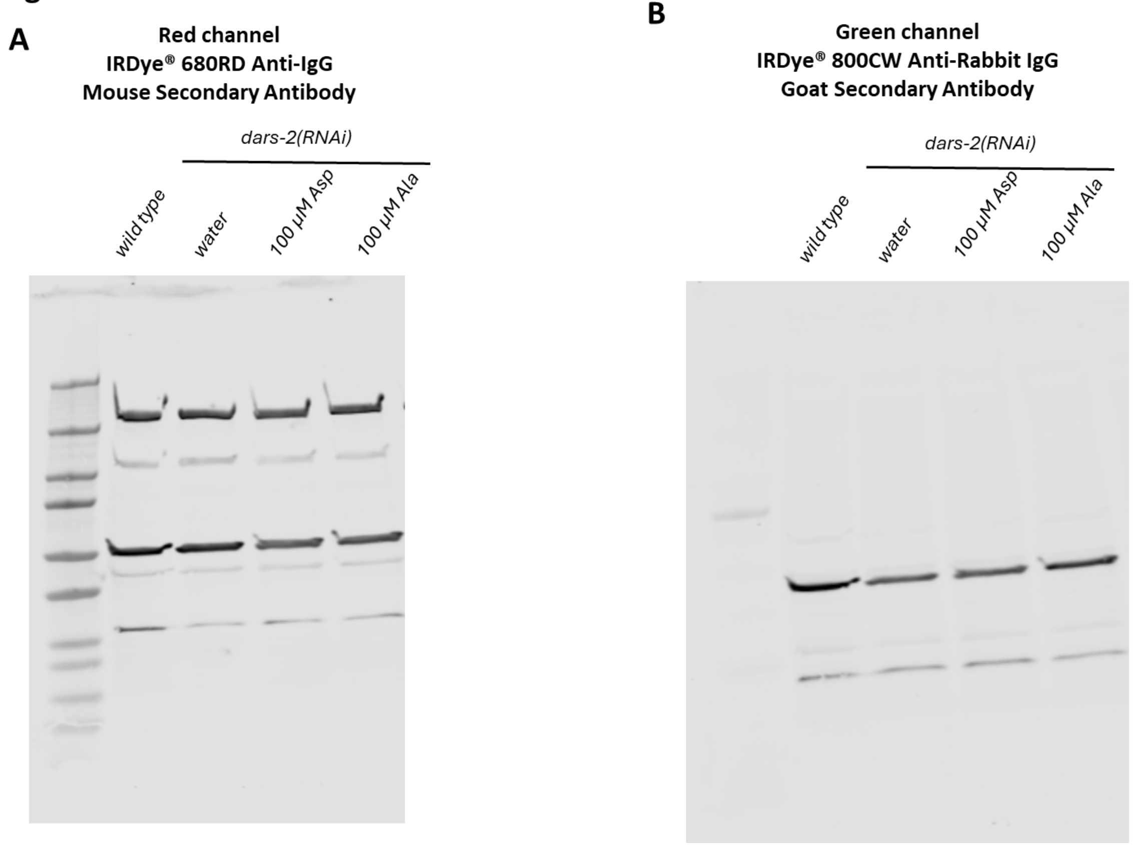


**Figure S6. Steady-state levels of OXPHOS proteins in *dars-2*(RNAi) worms following aspartate treatment. (A)** Full blot corresponding to Fig. 6A, visualized with the Total OXPHOS antibody in the red channel. **(B)** Full blot corresponding to Fig. 6A, visualized with Tom20 and β-actin antibodies in the green channel. **Related to Fig. 6.**

**Supplementary Table 1. Summary of RNAi plasmids used to knockdown individual mt-ARS genes in *C. elegans***

| Amino acid | Human gene | *C. elegans* gene | Source | *RNAi* clone | % amino acid similarity |
|---|---|---|---|---|---|
| Alanine | AARS2 - alanyl-tRNA synthetase 2 | *aars-1* | *C. elegans* ORFeome Library (Horizon Discovery) | W02B12.6 | 52% |
| Cysteine | CARS2 - cysteinyl-tRNA synthetase 2 | *cars-1* | *C. elegans* ORFeome Library (Horizon Discovery) | Y23H5A.7 | 36.7% |
| Aspartic acid | DARS2 - aspartyl-tRNA synthetase 2 | *dars-2* | *C. elegans* ORFeome Library (Horizon Discovery) | F10C2.6 | 56.4% |
| Glutamic acid | EARS2 - glutamyl-tRNA synthetase 2 | *ears-2* | *C. elegans* ORFeome Library (Horizon Discovery) | T07A9.2 | 56% |
| Phenylalanine | FARS2 - phenylalanyl-tRNA synthetase 2 | *fars-2* | C. elegans ORFeome Library (Horizon Discovery) | T08B2.9 | 62.8% |
| Glycine | GARS1 - glycyl-tRNA synthetase 1 | *gars-1* | C. elegans ORFeome Library (Horizon Discovery) | T10F2.1 | 57.7% |
| Histidine | HARS2 histidyl-tRNA synthetase 2 | *hars-1* | C. elegans ORFeome Library (Horizon Discovery) | T11G6.1 | 69.6% |
| Isoleucine | IARS2 - isoleucyl-tRNA synthetase 2 | *iars-2* | C. elegans ORFeome Library (Horizon Discovery) | C25A1.7 | 48.9% |
| Lysine | KARS1 - lysyl-tRNA synthetase 1, mitochondrial | *kars-1* | C. elegans ORFeome Library (Horizon Discovery) | T02G5.9 | 75.2% |
| Leucine | LARS2 - leucyl-tRNA synthetase 2 | *lars-2* | C. elegans ORFeome Library (Horizon Discovery) | *ZK524.3* | 52.3% |
| Methionine | MARS2 - methionyl-tRNA synthetase 2 | *mars-2* | C. elegans ORFeome Library (Horizon Discovery) | *Y105E8A.20* | 56.2% |
| Asparagine | NARS2 - asparaginyl-tRNA synthetase 2 | *nars-2* | C. elegans ORFeome Library (Horizon Discovery) | Y66D12A.23 | 52.3% |
| Proline | PARS2 prolyl-tRNA synthetase 2 | *pars-2* | C. elegans ORFeome Library (Horizon Discovery) | *T27F6.5* | 51.4% |
| Arginine | RARS2 arginyl-tRNA synthetase 2 | *rars-2* | C. elegans ORFeome Library (Horizon Discovery) | *C29H12.1* | 49.9% |
| Serine | SARS2- seryl-tRNA synthetase 2 | *sars-2* | Ahringer *C. elegans RNAi* feeding library (DNAFORM) | *W03B1.4* | 50.5% |

| | | | | | |
|---|---|---|---|---|---|
| Threonine | TARS2 - threonyl-tRNA synthetase 2 | *tars-1* | Ahringer *C. elegans RNAi* feeding library (DNAFORM) | *C47D12.6* | 48.3% |
| Valine | VARS2 - valyl-tRNA synthetase 2, | *glp-4* | Ahringer *C. elegans RNAi* feeding library (DNAFORM) | *Y87G2A.5* | 59.7% |
| Tryptophane | WARS2 - tryptophanyl tRNA synthetase 2 | *wars-2* | Ahringer *C. elegans RNAi* feeding library (DNAFORM) | *C34E10.4* | 42.7% |
| Tyrosine | YARS2 - tyrosyl-tRNA synthetase 2 | *yars-2* | Ahringer *C. elegans RNAi* feeding library (DNAFORM) | *K08F11.4* | 52.3% |
