## Supplementary material for "Cognate amino acid therapies provide preclinical benefit in 19 *C. elegans* models of ARS2 deficiency": tables

**Table 1. Overview of the gross phenotype changes observed following two consecutive generations of mt-ARS knockdown in worms.**

| mtARS Gene knocked down by RNAi | | First Generation *(RNAi)* | | | Second Generation *$(RNAi)^2$* | | |
|---|---|---|---|---|---|---|---|
| | | length | $UPR^{mt}$ stress | activity | length | $UPR^{mt}$ stress | activity |
| 1 | *aars-2* | ⬇ | ns | ns | ⬇ | ⬆ | ⬇ |
| 2 | *cars-1* | ⬇ | ns | ns | **sterile** | | |
| 3 | *dars-2* | ⬇ | ⬆ | ⬇ | ⬇ | ⬆ | ⬇ |
| 4 | *ears-2* | ⬇ | ⬆ | ns | ⬇ | ⬆ | ⬇ |
| 5 | *fars-2* | ⬇ | ns | ns | **sterile** | | |
| 6 | *gars-1* | ⬇ | ns | ns | ⬇ | ⬆ | ⬇ |
| 7 | *hars-2* | ⬇ | ns | ns | **sterile** | | |
| 8 | *iars-2* | ⬇ | ns | ns | ⬇ | ⬆ | ⬇ |
| 9 | *kars-1* | ⬇ | ⬆ | ns | **sterile** | | |
| 10 | *lars-2* | ⬇ | ns | ns | ⬇ | ⬆ | ns |
| 11 | *mars-2* | ⬇ | ⬆ | ns | **sterile** | | |
| 12 | *nars-2* | ⬇ | ns | ns | ⬇ | ⬆ | ⬇ |
| 13 | *pars-2* | ⬇ | ns | ns | ⬇ | ⬆ | ⬇ |
| 14 | *rars-2* | ⬇ | ns | ns | ⬇ | ⬆ | ns |
| 15 | *sars-2* | ⬇ | ⬆ | ⬇ | ⬇ | ⬆ | ⬇ |
| 16 | *tars-2* | ⬇ | ns | ns | ⬇ | ⬆ | ⬇ |
| 17 | *glp-4 (vars-2)* | ⬇ | ns | ⬇ | ⬇ | ⬆ | ⬇ |
| 18 | *wars-2* | ⬇ | ns | ns | ⬇ | ⬆ | ns |
| 19 | *yars-2* | ⬇ | ns | ns | ⬇ | ⬆ | ⬇ |
